# Variational Bayesian parameter estimation techniques for the general linear model

**DOI:** 10.1101/077461

**Authors:** Ludger Starke, Dirk Ostwald

## Abstract

Variational Bayes (VB), variational maximum likelihood (VML), restricted maximum likelihood (ReML), and maximum likelihood (ML) are cornerstone parametric statistical estimation techniques in the analysis of functional neuroimaging data. However, the theoretical underpinnings of these model parameter estimation techniques are rarely covered in introductory statistical texts. Because of the widespread practical use of VB, VML, ReML, and ML in the neuroimaging community, we reasoned that. a theoretical treatment of their relationships and their application in a basic modelling scenario may be helpful for both neuroimaging novices and practitioners alike. In this technical study, we thus revisit the conceptual and formal underpinnings of VB, VML, ReML, and ML and provide a detailed account of their mathematical relationships and implementational details. We further apply VB, VML, ReML, and ML to the general linear model (GLM) with non-spherical error covariance as commonly encountered in the first-level analysis of fMRI data. To this end, we explicitly derive the corresponding free energy objective functions and ensuing iterative algorithms. Finally, in the applied part of our study, we evaluate the parameter and model recovery properties of VB, VML, ReML, and ML, first in an exemplary setting and then in the analysis of experimental fMRI data acquired from a single participant under visual stimulation.

## 1 Introduction

Variational Bayes (VB), variational maximum likelihood (VML) (also known as expectation-maximization), restricted maximum likelihood (ReML), and maximum likelihood (ML) are cornerstone parametric statistical estimation techniques in the analysis of functional neuroimaging data. In the SPM software environment (http://www.fil.ion.ucl.ac.uk/spm/), one of the most commonly used software packages in the neuroimaging community, variants of these estimation techniques have been implemented for a wide range of data models (Ashburner, 2012; Penny et al., 2011). For fMRI data, these models vary from mass-univariate general linear and auto-regressive models (e.g., Friston et al., 1994, 2002a,b; Penny et al., 2003), over multivariate decoding models (e.g., Friston et al., 2008a), to dynamic causal models (e.g., Friston et al., 2003; Stephan et al., 2008; Marreiros et al., 2008). For M/EEG data, these models range from channel-space general linear models (e.g., Kiebel and Friston, 2004a,b), over dipole and distributed source reconstruction models (e.g., Kiebel et al., 2008; Friston et al., 2008b; Litvak and Friston, 2008), to a large family of dynamic causal models (e.g., David et al., 2006; Chen et al., 2008; Moran et al., 2009; Pinotsis et al., 2012; Ostwald and Starke, 2016).

Because VB, VML, ReML, and ML determine the scientific inferences drawn from empirical data in any of the above mentioned modelling frameworks, they are of immense importance for the neuroimaging practitioner. However, the theoretical underpinnings of these estimation techniques are rarely covered in introductory statistical texts and the technical literature relating to these techniques is rather evolved. Because of their widespread use within the neuroimaging community, we reasoned that a theoretical treatment of these techniques in a familiar model scenario may be helpful for both neuroimaging novices, who would like to learn about some of the standard statistical estimation techniques employed in the field, and for neuroimaging practitioners, who would like to further explore the foundations of these and alternative model estimation approaches.

In this technical study, we thus revisit the conceptual underpinnings of the aforementioned techniques and provide a detailed account of their mathematical relations and implementational details. Our exposition is guided by the fundamental insight that VML, ReML, and ML can be understood as special cases of VB (Friston et al., 2002a, 2007; Friston, 2008). In the current note, we reiterate and consolidate this conceptualization by paying particular attention to the respective technique’s formal treatment of a model’s parameter set. Specifically, across the estimation techniques of interest, model parameters are either treated as random variables, in which case they are endowed with prior and posterior uncertainty modelled by parametric probability density functions, or as non-random quantities. In the latter case, prior and posterior uncertainties about the respective parameters’ values are left unspecified. Because the focus of the current account is on statistical estimation techniques, we restrict the model of application to a very basic scenario that every neuroimaging practitioner is familiar with: the analysis of a single-participant, single-session EPI time-series in the framework of the general linear model (GLM) (Monti, 2011; Poline and Brett, 2012). Importantly, in line with the standard practice in fMRI data analysis, we do not assume spherical covariance matrices (e.g., Mumford and Nichols, 2008; Zarahn et al., 1997; Purdon and Weisskoff, 1998; Woolrich et al., 2001; Friston et al., 2002b).

We proceed as follows. After some preliminary notational remarks, we begin the theoretical exposition by first introducing the model of application in Section 2.1. We next briefly discuss two standard estimation techniques (conjugate Bayes and ML for spherical covariance matrices) that effectively span the space of VB, VML, ReML, and ML and serve as useful reference points in Section 2.2. After this prelude, we are then concerned with the central estimation techniques of interest herein. In a hierarchical fashion, we subsequently discuss the theoretical background and the practical algorithmic application of VB, VML, ReML, and ML to the GLM in Sections 2.3 - 2.6. We focus on the central aspects and conceptual relationships of the techniques and present all mathematical derivations as Supplementary Material. In the applied part of our study (Section 3), we then firstly evaluate VB, VML, ReML, and ML from an objective Bayesian viewpoint (Bernardo, 2009) in simulations; and secondly, apply them to real fMRI data acquired from a single participant under visual stimulation (Ostwald et al., 2010). We close by discussing the relevance and relation of our exposition with respect to previous treatments of the topic matter in Section 4.

In summary, we make the following novel contributions in the current technical study. Firstly, we provide a comprehensive mathematical documentation and derivation of the conceptual relationships between VB, VML, ReML, and ML. Secondly, we derive a collection of explicit algorithms for the application of these estimation techniques to the GLM with non-spherical linearized covariance matrix. Finally, we explore the validity of the ensuing algorithms in simulations and in the application to real experimental fMRI data. We complement our theoretical documentation by the practical implementation of the algorithms and simulations in a collection of Matlab .m files (MATLAB and Optimization Toolbox Release 2014b, The MathWorks, Inc., Natick, MA, United States), which is available from the Open Science Framework (https://osf.io/c4ux7/). On occasion, we make explicit reference to these functions, which share the stub *vbg_*.m*.

### Notation and preliminary remarks

A few remarks on our mathematical notation are in order. We formulate VB, VML, ReML, and ML against the background of probabilistic models (e.g., Bishop, 2006; Barber, 2012; Murphy, 2012). By probabilistic models we understand (joint) probability distributions over sets of observed and unobserved random variables. Notationally, we do not distinguish between probability distributions and their associated probability density functions and write, for example, *p*(*y, θ*) for both. Because we are only concerned with parametric probabilistic models of the Gaussian type, we assume throughout the main text that all probability distributions of real random vectors have densities. We do, however, distinguish between the conditioning of a probability distribution of a random variable *y* on a (commonly unobserved) random variable *θ*, which we denote by *p*(*y|θ*), and the parameterization of a probability distribution of a random variable *y* by a (non-random) parameter *θ*, which we denote by *p_θ_*(*y*). Importantly, in the former case, *θ* is conceived of as random variable, while in the latter case, it is not. Equivalently, if *θ*\* denotes a value that the random variable *θ* may take on, we set 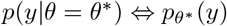.

Otherwise, we use standard applied mathematical notation. For example, real vectors and matrices are denoted as elements of 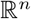 and 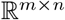 for 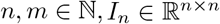 denotes the *n*-dimensional identity matrix, | · | denotes a matrix determinant, tr(·) denotes the trace operator, and p.d. denotes a positive-definite matrix. *H_f_* (*a*) denotes the Hessian matrix of some real-valued function *f* (*x*) evaluated at *x* = *a*. We denote the probability density function of a Gaussian distributed random vector *y* with expectation parameter *μ* and covariance parameter Σ by *N*(*y*; *μ*, Σ). Finally, because of the rather applied character of this note, we formulate functions primarily by means of the definition of the values they take on and eschew formal definitions of their domains and ranges. Further notational conventions that apply in the context of the mathematical derivations provided in the Supplementary Material are provided therein.

## 2 Theory

### 2.1 Model of interest

Throughout this study, we are interested in estimating the parameters of the model

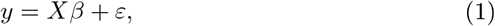

where 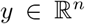 denotes the data, 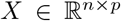 denotes a design matrix of full column rank *p*, and 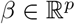 denotes a parameter vector. We make the following fundamental assumption about the error term 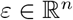

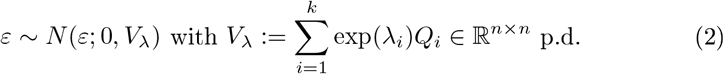

In words, we assume that the error term is distributed according to a Gaussian distribution with expectation parameter 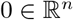 and positive-definite covariance matrix 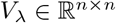. Importantly, we do not assume that *V_λ_* is necessarily of the form *σ*^2^*I_n_*, i.e. we allow for non-sphericity of the error terms. In (2), *λ*_1_, …, *λ_k_*, is a set of *covariance component parameters* and 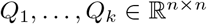 is a set of *covariance basis matrices*, which are assumed to be fixed and known. We assume throughout, that the true, but unknown, values of *λ*_1_, …, *λ_k_* are such that *V_λ_* is positive-definite. In line with the common denotation in the neuroimaging literature, we refer to (1) and (2) as the *general linear model* (GLM) and its formulation by means of equations (1) and (2) as its *structural form*.

Models of the form (1) and (2) are widely used in the analysis of neuroimaging data, and, in fact, throughout the empirical sciences (e.g., Rutherford, 2001; Draper and Smith, 2014; Gelman et al., 2014). In the neuroimaging community, models of the form (1) and (2) are used, for example, in the analysis of fMRI voxel time-series at the session and participant-level (Monti, 2011; Poline and Brett, 2012), for the analysis of group effects (Mumford and Nichols, 2006, 2009), or in the context of voxel-based morphometry (Ashburner and Friston, 2000; Ashburner, 2009).

In the following, we discuss the application of VB, VML, ReML, and ML to the general forms of (1) and (2). In our examples, however, we limit ourselves to the application of the GLM in the analysis of a single voxel’s time-series in a single fMRI recording (run). In this case, 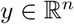 corresponds to the voxel’s MR values over EPI volume acquisitions and 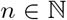 represents the total number of volumes acquired during the session. The design matrix 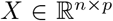 commonly constitutes a constant regressor and the onset stick functions of different experimental conditions convolved with a haemodynamic response function and a constant offset. This renders the parameter entries 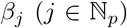 to correspond to the average session MR signal and condition-specific effects. Importantly, in the context of fMRI time-series analyses, the most commonly used form of the covariance matrix *V_λ_* employs *k* = 2 covariance component parameters *λ*_1_ and *λ*_2_ and corresponding covariance basis matrices

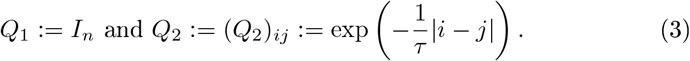

This specific form of the error covariance matrix encodes exponentially decaying correlations between neighbouring data points, and, with *τ* := 0.2, corresponds to the widely used approximation to the *AR(1) + white noise* model in the analysis of fMRI data (Purdon and Weisskoff, 1998; Friston et al., 2002b).

In Figure 1, we visualize the exemplary design matrix and covariance basis matrix set that will be employed in the example applications throughout the current section. In the example, we assume two experimental conditions, which have been presented with an expected inter-trial interval of 6 seconds (standard deviation 1 second) during an fMRI recording session comprising *n* = 400 volumes and with a TR of 2 seconds. The design matrix was created using the micro-time resolution convolution and downsampling approach discussed in Henson and Friston (2007).

**Figure 1:**
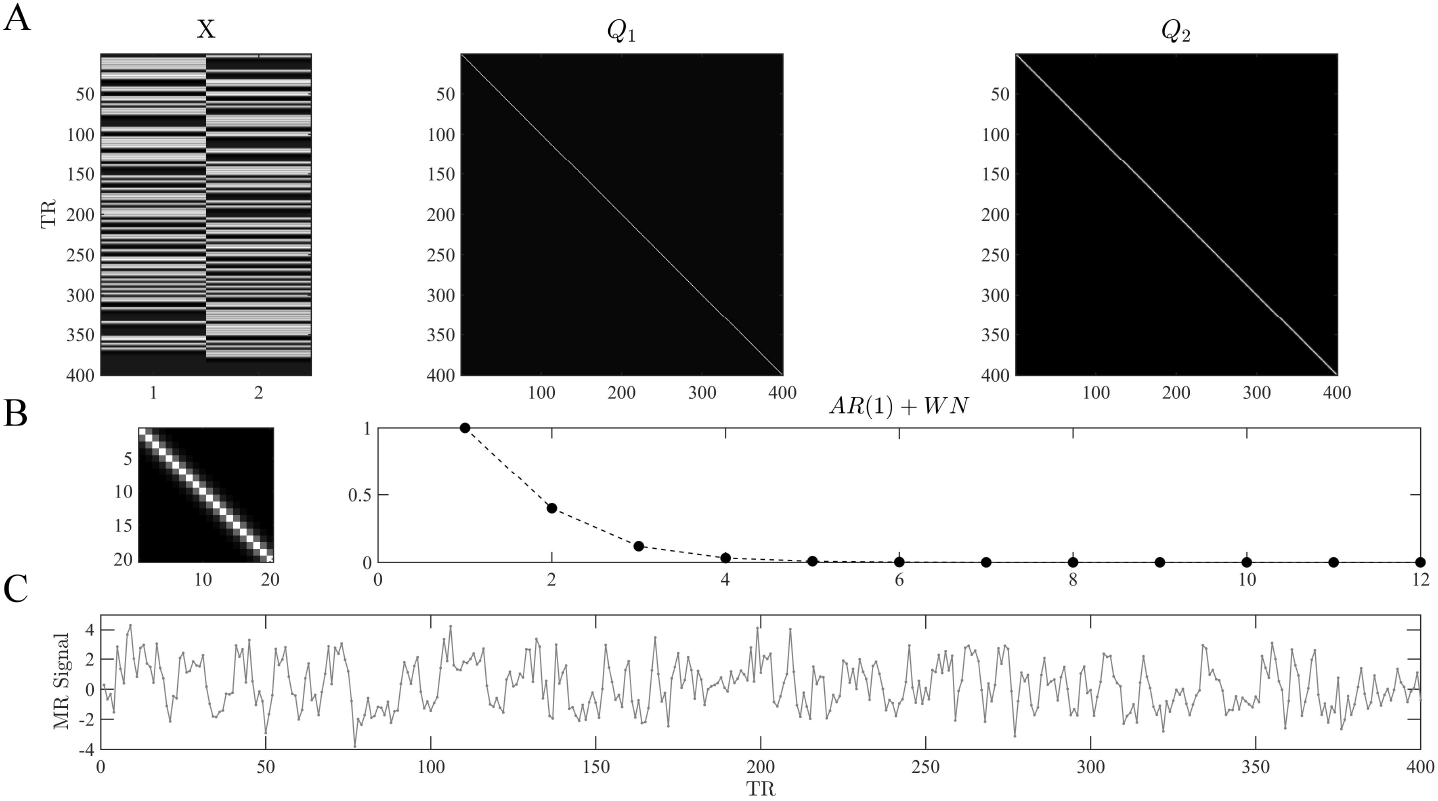
(A) Example design and covariance basis matrices. The upper panels depict the design matrix 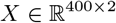 and the covariance basis matrices 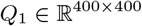 used in the example applications of the current section. The design matrix encodes the onset functions of two hypothetical experimental conditions which were convolved with the canonical haemodynamic response function. Events of each condition are presented approximately every 6 seconds, and *n* = 400 data points with a TR of 2 seconds are modelled. The covariance basis matrices are specified in eq.(3) and shown here for *n* = 400 based on their evaluation using *spm_Ce.m*. (B) The left panel depicts a magnification of the first 20 entries of *Q*_2_. The right panel depicts the entries of the first row of *Q*_2_ for 12 columns. For *τ* = 0.2 the entries model exponentially decaying error correlations. (C) A data realization of the ensuing GLM model with true, but unknown, values of *β* = (2, −1)^*T*^ and *λ* = (−0.5, −2)^*T*^. Note that we do not model a signal offset, or equivalently, set the beta parameter for the signal offset to zero. For implementational details, please see *vbg_1.m*.

### 2.2 Conjugate Bayes and ML under error sphericity

We start by briefly recalling the fundamental results of conjugate Bayesian and classical point-estimation for the GLM with spherical error covariance matrix. In fact, the introduction of ReML (Phillips et al., 2002; Friston et al., 2002a) and later VB (Friston et al., 2007) to the neuroimaging literature were motivated amongst other things by the need to account for non-sphericity of the error distributions in fMRI time-series analysis (Purdon and Weisskoff, 1998; Woolrich et al., 2001). Further, while not a common approach in fMRI, recalling the conjugate Bayes scenario helps to contrast the probabilistic model of interest in VB from its mathematically more tractable, but perhaps less intuitively plausible, analytical counterpart. Together, the two estimation techniques discussed in the current section may thus be conceived as forming the respective endpoints of the continuum of estimation techniques discussed in the remainder.

With spherical covariance matrix, the GLM of eqs. (1) and (2) simplifies to

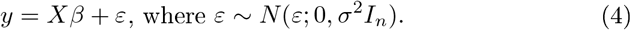

A conjugate Bayesian treatment of the GLM considers the structural form (4) as a conditional probabilistic statement about the distribution of the observed random variable *y*

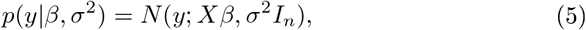

which is referred to as the *likelihood* and requires the specification of the marginal distribution *p*(*β, σ*^2^), referred to as the *prior*. Together, the likelihood and the prior define the probabilistic model of interest, which takes the form of a joint distribution over the observed random variable *y* and the unobserved random variables *β* and *σ*^2^:

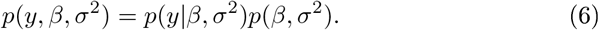

Based on the probabilistic model (6), the two fundamental aims of Bayesian inference are, firstly, to determine the conditional parameter distribution given a value of the observed random variable *p*(*β, σ*^2^|*y*), often referred to as the *posterior*, and secondly, to evaluate the marginal probability *p*(*y*) of a value of the observed random variable, often referred to as *marginal likelihood* or *model evidence*. The latter quantity forms an essential precursor for Bayesian model comparison, as discussed for example in further detail in Stephan et al. (2016a). Note that in our treatment of the Bayesian scenario the marginal and conditional probability distributions of *β* and *σ*^2^ are meant to capture our uncertainty about the values of these parameters and not distributions of true, but unknown, parameter values. For the true, but unknown, values of *β* and *σ*^2^ we postulate, as in the classical point-estimation scenario, that they assume fixed values, which are never revealed (but can of course be chosen ad libitum in simulations).

The VB treatment of (6) assumes proper prior distributions for *β* and *σ*^2^. In this spirit, the closest conjugate Bayesian equivalent is hence the assumption of proper prior distributions. For the case of the model (6), upon reparameterization in terms of a precision parameter *λ* := 1/*σ*^2^, a natural conjugate approach assumes a non-independent prior distribution of Gaussian-Gamma form,

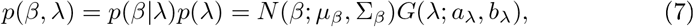

where 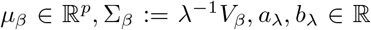 are the prior distribution parameters and 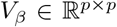 p.d. is the prior beta parameter covariance structure. For the gamma distribution we use the shape and rate parameterization. Notably, the Gaussian distribution of *β* is parameterized conditional on the value of *λ* in terms of its covariance Σ*_β_*. Under this prior assumption, it can be shown that the posterior distribution is also of Gaussian-Gamma form,

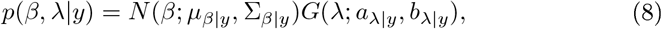

with posterior parameters

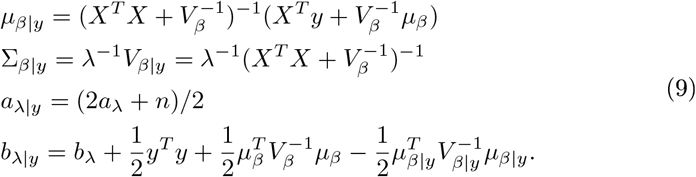

Furthermore, in this scenario the marginal likelihood evaluates to a multivariate non-central T-distribution

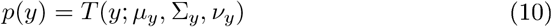

with expectation, covariance, and degrees of freedom parameters

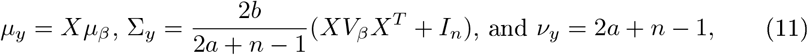

respectively. For derivations of (8) - (11) see, for example, Lindley and Smith (1972); Broemeling (1984), and Gelman et al. (2014).

Importantly, in contrast to the VB, VML, ReML, and ML estimation techniques developed in the remainder, the assumption of the prior probabilistic dependency of the effect size parameter on the covariance component parameter in (7) eshews the need for iterative approaches and results in the fully analytical solutions of eqs. (8) to (11). However, as there is no principled reason beyond mathematical convenience that motivates this prior dependency, the fully conjugate framework seems to be rarely used in the analysis of neuroimaging data. Moreover, the assumption of an uninformative improper prior distribution (Frank et al., 1998) is likely more prevalent in the neuromaging community than the natural conjugate form discussed above. This is due to the implementation of a closely related procedure in FSL’s FLAME software (Woolrich et al., 2004, 2009). However, because VB assumes proper prior distributions, we eschew the details of this approach herein.

In contrast to the probabilistic model of the Bayesian scenario, the classical ML approach for the GLM does not conceive of *β* and *σ*^2^ as unobserved random variables, but as parameters, for which point-estimates are desired. The probabilistic model of the classical ML approach for the structural model (4) thus takes the form

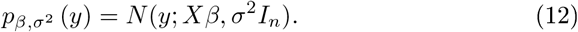

The ML point-estimators for *β* and *σ*^2^ are well-known to evaluate to (e.g., Hocking, 2013).

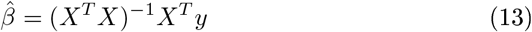

and

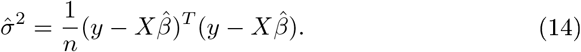

Note that (13) also corresponds to the ordinary least-squares estimator. It can be readily generalized for non-spherical error covariance matrices by a “sandwiched” inclusion of the appropriate error covariance matrix, if this is (assumed) to be known, resulting in the generalized least-squares estimator (e.g., Draper and Smith, 2014). Further note that (14) is a biased estimator for *σ*^2^ and hence commonly replaced by its restricted maximum likelihood counterpart, which replaces the factor *n*^−1^ by the factor (*n* − *p*)^−1^ (e.g., Foulley, 1993).

Having briefly reviewed the conjugate Bayesian and classical point estimation techniques for the GLM parameters under the assumption of a spherical error covariance matrix, we next discuss VB, VML, ReML, and ML for the scenario laid out in Section 2.1.

### 2.3 Variational Bayes (VB)

VB is a computational technique that allows for the evaluation of the primary quantities of interest in the Bayesian paradigm as introduced above: the posterior parameter distribution and the marginal likelihood. For the GLM, VB thus rests on the same probabilistic model as standard conjugate Bayesian inference: the structural form of the GLM (cf. equations (1) and (2)) is understood as the parameter conditional likelihood distribution and both parameters are endowed with marginal distributions. The probabilistic model of interest in VB thus takes the form

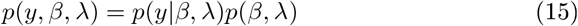

with likelihood distribution

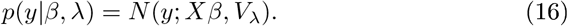

Above, we have seen that a conjugate prior distribution can be constructed which allows for exact inference in models of the form (1) and (2) based on a conditionally-dependent prior distribution and simple covariance form. In order to motivate the application of the VB technique to the GLM, we here thus assume that the marginal distribution *p*(*β, λ*) factorizes, i.e., that

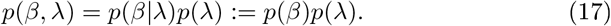

Under this assumption, exact Bayesian inference for the GLM is no longer possible and approximate Bayesian inference is clearly motivated (Murphy, 2012).

To compute the marginal likelihood and obtain an approximation to the posterior distribution over parameters *p*(*β, λ|y*), VB uses the following decomposition of the log marginal likelihood into two information theoretic quantities (Cover and Thomas, 2012), the *free energy* and a *Kullback-Leibler (KL) divergence*

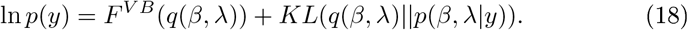

We discuss the constituents of the right-hand side of (18) in turn. Firstly, *q*(*β, λ*) denotes the so-called *variational distribution*, which will constitute the approximation to the posterior distribution and is of parameterized form, i.e. governed by a probability density. We refer to the parameters of the variational distribution as *variational parameters*. Secondly, the non-negative KL-divergence is defined as the integral

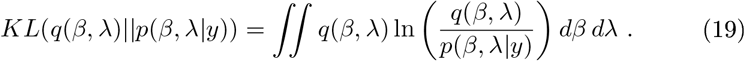

Note that, formally, the KL-divergence is a functional, i.e., a function of functions, in this case the probability density functions *q*(*β, λ*) and *p*(*β, λ|y*), and returns a scalar number. Intuitively, it measures the dissimilarity between its two input distributions: the more similar the variational distribution *q*(*β, λ*) is to the posterior distribution *p*(*β, λ|y*), the smaller the divergence becomes. It is of fundamental importance for the VB technique that the KL-divergence is always positive and zero if, and only if, *q*(*β, λ*) and *p*(*β, λ|y*) are equal. For a proof of these properties, see Appendix A in Ostwald et al. (2014). Together with the log marginal likelihood decomposition (18) the properties of the KL-divergence equip the free energy with its central properties for the VB technique, as discussed below. A proof of (18) with *ϑ* := {*β, λ*} is provided in Appendix B in Ostwald et al. (2014).

The free energy itself is defined by

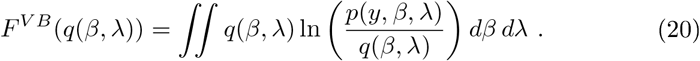

Due to the non-negativity of the KL-divergence, the free energy is always smaller than or equal to the log marginal likelihood - the free energy thus forms a lower bound to the log marginal likelihood. Note that in (20), the data *y* is assumed to be fixed, such that the free energy is a function of the variational distribution only. Because, for a given data observation, the log marginal likelihood ln *p*(*y*) is a fixed quantity, and because increasing the free energy contribution to the right-hand side of (18) necessarily decreases the KL-divergence between the variational and the true posterior distribution, maximization of the free energy with respect to the variational distribution has two consequences: firstly, it renders the free energy an increasingly better approximation to the log marginal likelihood; secondly, it renders the variational approximation an increasingly better approximation to the posterior distribution.

In summary, VB rests on finding a variational distribution that is as similar as possible to the posterior distribution, which is equivalent to maximizing the free energy with regard to the variational distribution. The maximized free energy then substitutes for the log marginal likelihood and the corresponding variational distribution yields an approximation to the posterior parameter distribution, i.e.,

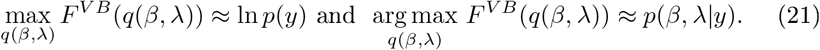

To facilitate the maximization process, the variational distribution is often assumed to factorize over parameter sets, an assumption commonly referred to as *mean-field approximation* (Friston et al., 2007)

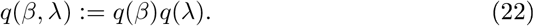

Of course, if the posterior does not factorize accordingly, i.e., if

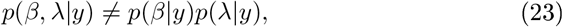

the mean-field approximation limits the exactness of the method.

In applications, maximization of the free energy is commonly achieved by either *free-form* or *fixed-form* schemes. In brief, free-form maximization schemes do not assume a specific form of the variational distribution, but employ a fundamental theorem of variational calculus to maximize the free energy and to analytically derive the functional form and parameters of the variational distribution. For more general features of the free-form approach, please see, for example, Bishop (2006); Chappell et al. (2009) and Ostwald et al. (2014). Fixed-form maximization schemes, on the other hand, assume a specific parametric form for the variational distribution’s probability density function from the outset. Under this assumption, the free energy integral (20) can be evaluated (or at least approximated) analytically and rendered a function of the variational parameters. This function can in turn be optimized using standard nonlinear optimization algorithms. In the following section, we apply a fixed-form VB approach to the current model of interest.

#### Application to the GLM

To demonstrate the fixed-form VB approach to the GLM of eqs. (1) and (2), we need to specify the parametric forms of the prior distributions *p*(*β*) and *p*(*λ*), as well as the parametric forms of the variational distribution factors *q*(*β*) and *q*(*λ*). Here, we assume that all these marginal distributions are Gaussian, and hence specified in terms of their expectation and covariance parameters:

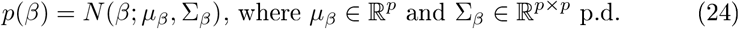

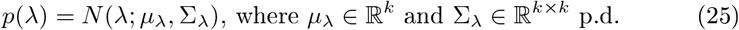

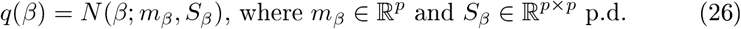

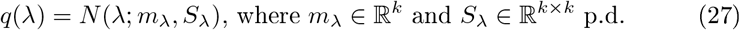

Note that we denote parameters of the prior distributions with Greek and parameters of the variational distributions with Roman letters. Together with eqs. (1) to (3), eqs. (24) to (27) specify all distributions necessary to evaluate the free energy integral and render the free energy a function of the variational parameters. We document this derivation in Supplementary Material S1.2 and here limit ourselves to the presentation of the result: under the given assumptions about the prior, likelihood, and variational distributions, the variational free energy is a function of the variational parameters *m_β_, S_β_, m_λ_*, and *S_λ_*, and, using mild approximations in its analytical derivation, evaluates to

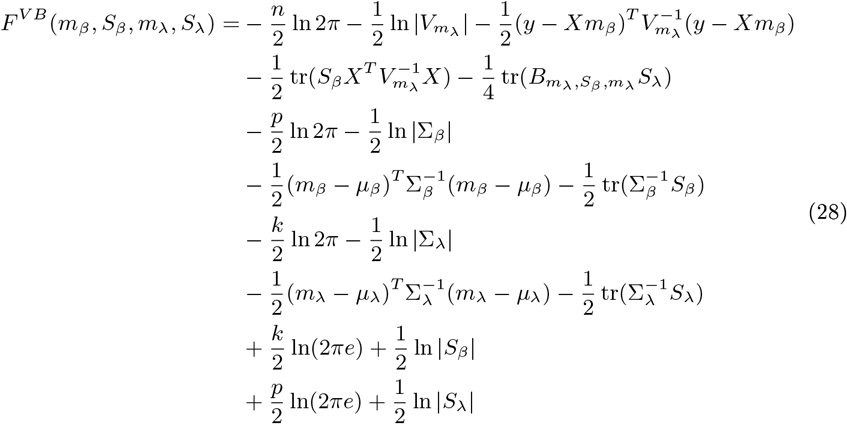

with

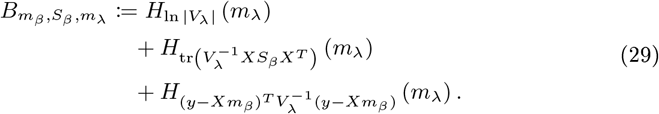

In (28), the third term may be viewed as an *accuracy term* which measures the deviation of the estimated model prediction from the data, the eighth and twelfth terms may be viewed as *complexity terms*, that measure how far the model can and has to deviate from its prior expectations to account for the data, and the last four terms can be conceived as *maximum entropy* terms that ensure that the posterior parameter uncertainty is as large as possible given the available data (Jaynes, 2003).

In principle, any numerical routine for the maximization of nonlinear functions could be applied to maximize the free energy function of eq. (28) with respect to its parameters. Because of the relative simplicity of eq. (28), we derived explicit update equations by evaluating the VB free energy gradient with respect to each of the parameters and setting to zero as documented in Supplementary Material S1.2. This analytical approach yields a set of four update equations and, together with the iterative evaluation of the VB free energy function (28), results in a VB algorithm for the current model as documented in Algorithm 1. Here, and in all remaining algorithms, convergence is assessed in terms of a vanishing of the free energy increase between successive iterations. This difference is evaluated against a convergence criterion *δ*, which we set to *δ* = 10^−3^ for all reported simulations.

In Figure 2, we visualize the application of the VB algorithm to an example fMRI time-series realization from the model described in Section 2.1 with true, but unknown, parameter values *β* = (2, −1)^*T*^ and *λ* = (−0.5, −2)^*T*^. We used imprecise priors for both *β* and *λ* by setting

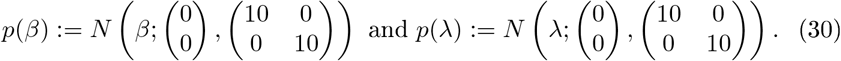

**Figure 2:**
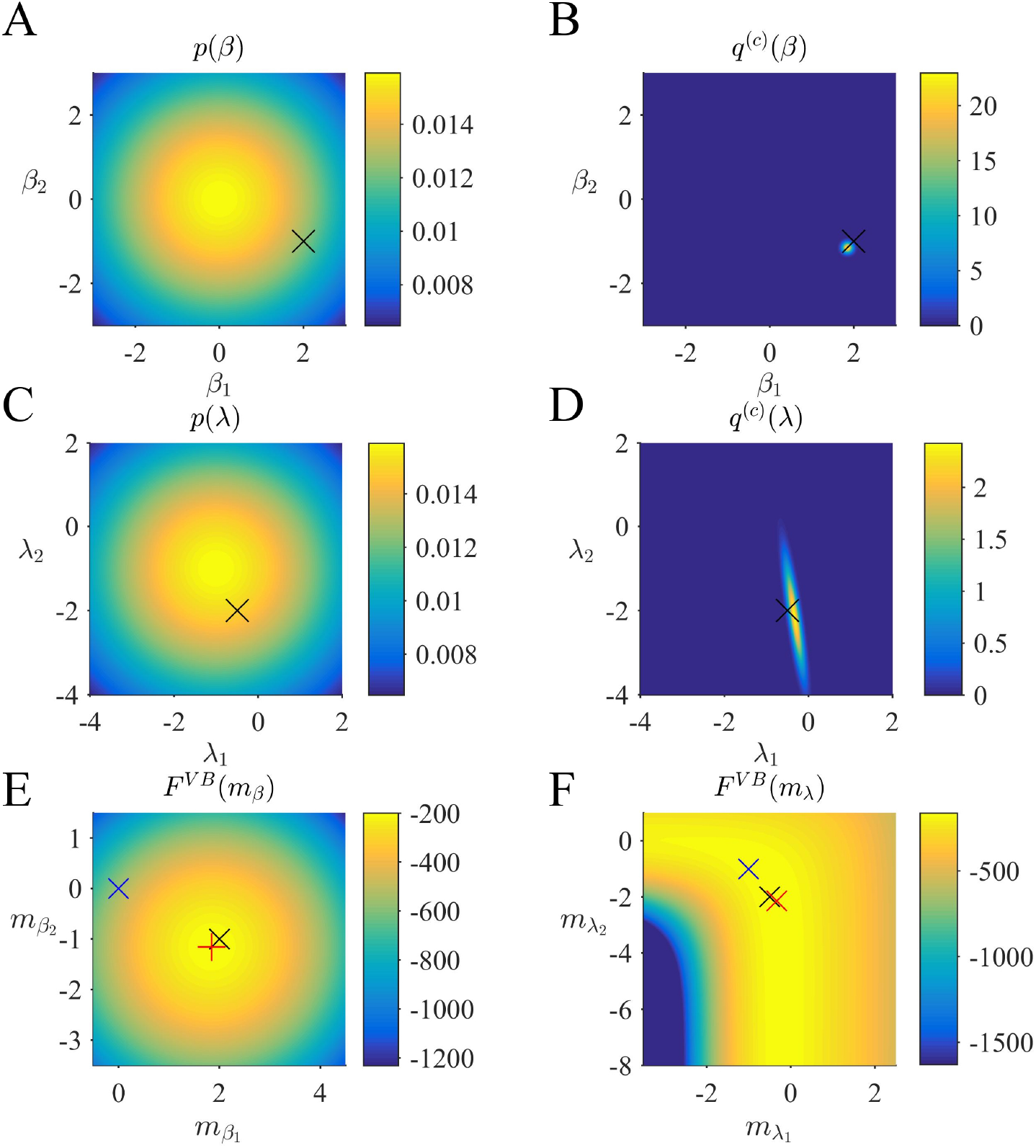
VB estimation. (A) Prior distribution *p*(*β*) with expectation *μ_β_* := (0, 0)^*T*^ and covariance Σ_*β*_ := 10*I*_2_. Here, and in all subpanels, the black x marks the true, but unknown, parameter value. (B) Variational approximation *q*^(*c*)^ (*β*) to the posterior distribution upon convergence (*δ* = 10^−3^). (C) Prior distribution *p*(*λ*) with expectation *μ_λ_* := (0, 0)^*T*^ and covariance Σ*_λ_* = 10*I*_2_. (D) Variational approximation *q*^(*c*)^ (*λ*) to the posterior distribution upon convergence. (E) Variational free energy dependence on *m_β_*. The blue x indicates the prior expectation parameter and the red + marks the approximated posterior expectation parameter. (F) Variational free energy dependence on *m_λ_*. The blue x indicates the prior expectation parameter and the red x details, please see *vbg_1.m*.

**Algorithm 1.**
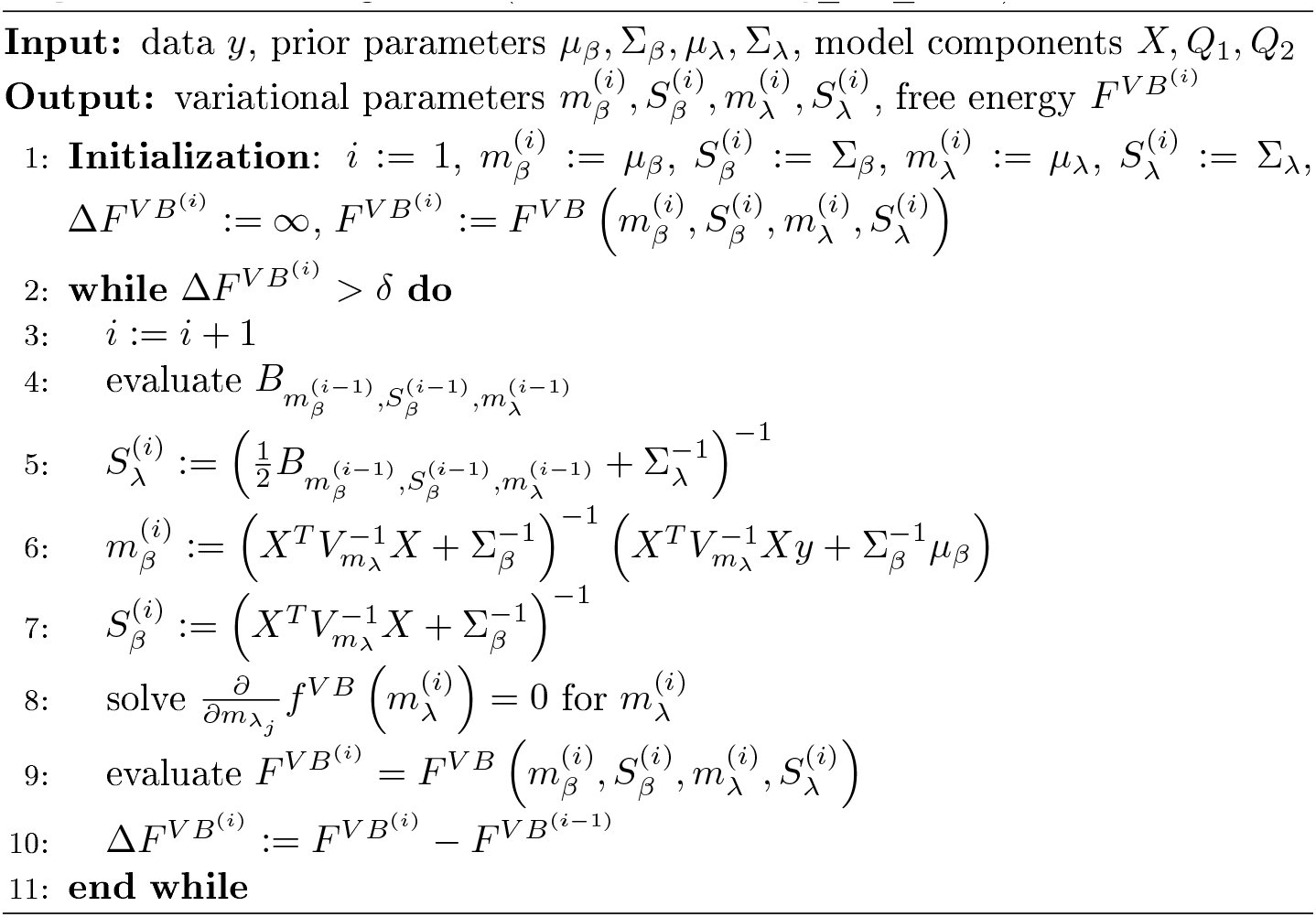
VB Algorithm (for details, see *vbg_est_vb.m*)

Panel A of Figure 2 depicts the prior distribution over *β*, and the true, but unknown, value of *β* as black ×. Panel B depicts the variational distribution over *β* after convergence for a VB free energy convergence criterion of *δ* = 10^−3^. Given the imprecise prior distribution, this variational distribution falls close to the true, but unknown, value. In general, convergence of the algorithm is achieved within 4 to 6 iterations. Panels C and D depict the prior distribution over *λ* and the variational distribution over *λ* upon convergence, respectively. As for *β*, the approximation of the posterior distribution is close to the true, but unknown, value of *λ*. Finally, Panels E and F depict the VB free energy surface as a function of the variational parameters *m_β_* and *m_λ_*, respectively. For the chosen prior distributions, the VB free energy surfaces display clear global maxima, which the VB algorithm can identify. Note, however, that the maximum of the VB free energy as a function of *m_λ_* is located on an elongated crest.

### 2.4 Variational Maximum Likelihood (VML)

Variational Maximum Likelihood (Beal, 2003), also referred to as (variational) expectation-maximization (Barber, 2012; McLachlan and Krishnan, 2007), can be considered a semi-Bayesian estimation approach. For a subset of model parameters, VML determines a Bayesian posterior distribution, while for the remaining parameters maximum-likelihood point estimates are evaluated. As discussed below, VML can be derived as a special case of VB under specific assumptions about the posterior distribution of the parameter set for which only point estimates are desired. If for this parameter set additionally a constant, improper prior is assumed, variational Bayesian inference directly yields VML estimates. In its application to the GLM, we here choose to treat *β* as the parameter for which a posterior distribution is derived, and *λ* as the parameter for which a point-estimate is desired.

The current probabilistic model of interest thus takes the form

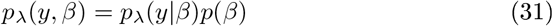

with likelihood distribution

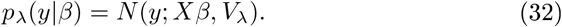

Note that in contrast to the probabilistic model underlying VB estimation, *λ* is not treated as a random variable and thus merely parameterizes the joint distribution of *β* and *y*. Similar to VB, VML rests on a decomposition of the log marginal likelihood

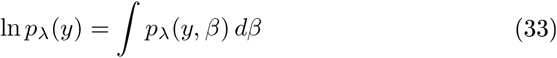

into a free energy and a KL-divergence term

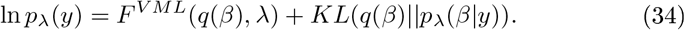

In contrast to the VB free energy, the VML free energy is defined by

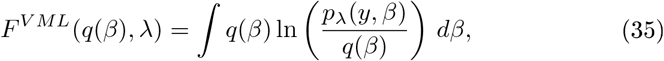

while the KL divergence term takes the form

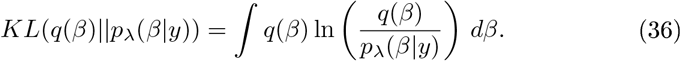

In Supplementary Material S2, we show how the VML framework can be derived as a special case of VB by assuming an improper prior for *λ* and a Dirac measure *δ*_*λ*\*_ for the variational distribution of *λ*. Importantly, it is the parameter value *λ*\* of the Dirac measure that corresponds to the parameter *λ* in the VML framework.

#### Application to the GLM

In the application of the VML approach to the GLM of eqs. (1) and (2) we need to specify the parametric forms of the prior distribution *p*(*β*) and the parametric form of the variational distribution *q*(*β*). As above, we assume that these distributions are Gaussian, i.e.,

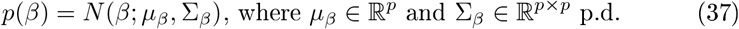

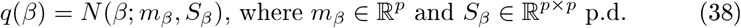

Based on the specifications of eqs. (37) and (38), the integral definition of the VML free energy can be analytically evaluated under mild approximations, which yields the VML free energy function of the variational parameters *m_β_* and *S_β_* and the parameter *λ*

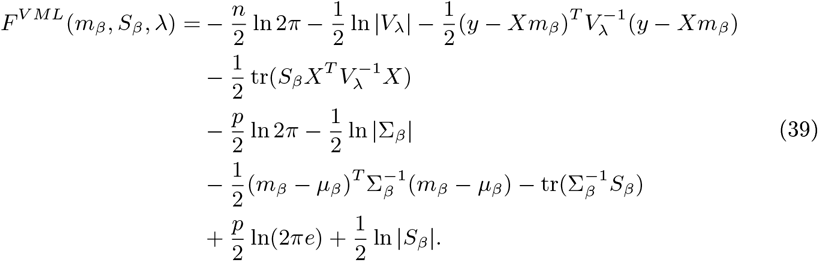

We document the derivation of (39) in Supplementary Material S1.3. In contrast to the VB free energy (cf. eq. (28)), the VML free energy for the GLM is characterized by the absence of terms relating to the prior and posterior uncertainty about the covariance component parameter *λ*. To maximize the VML free energy, we again derived a set of update equations as documented in Supplementary Material S1.3. These update equations give rise to a VML algorithm for the current model, which we document in Algorithm 2.

**Algorithm 2.**
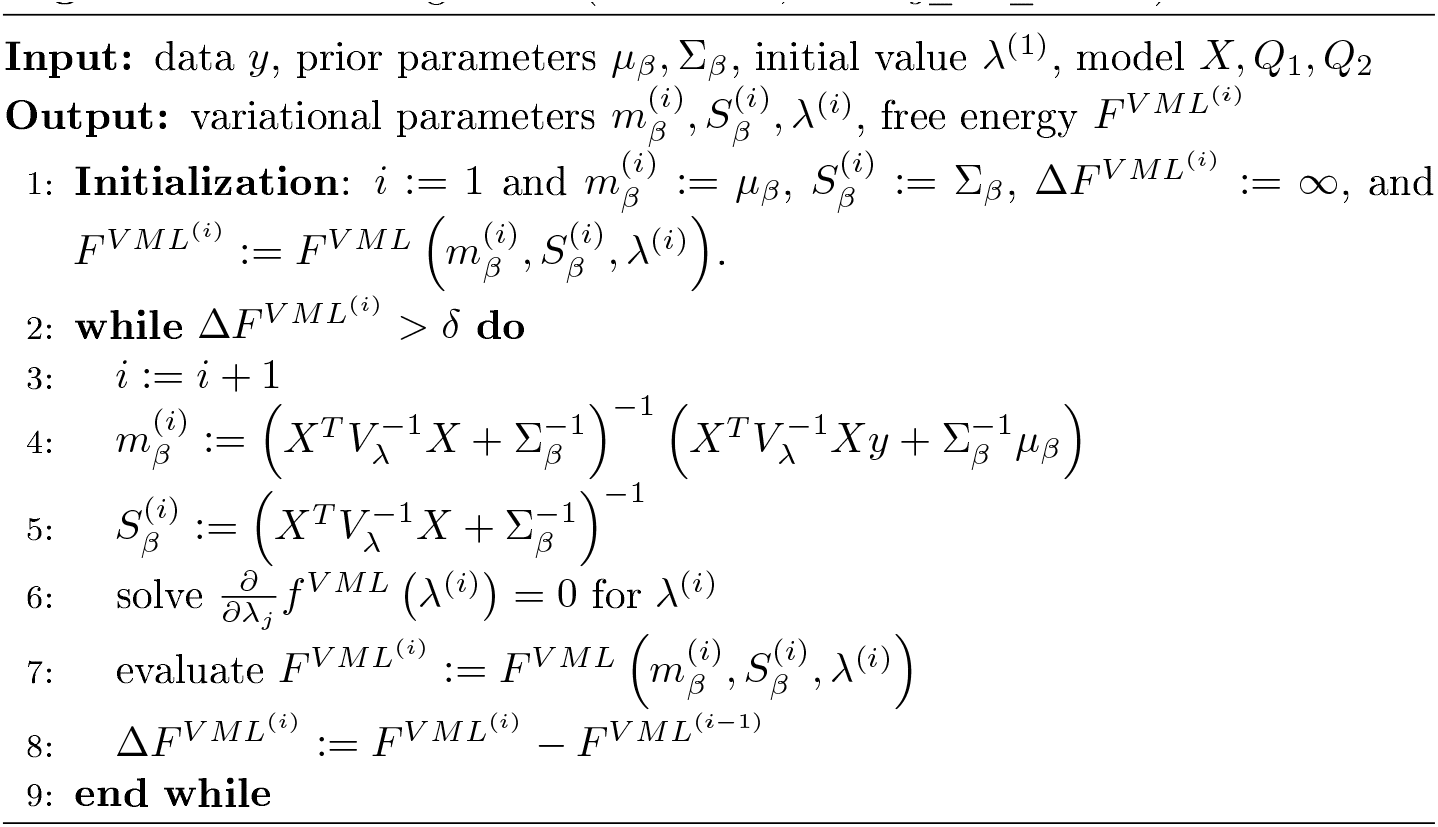
VML Algorithm (for details, see *vbg_est_vml.m*)

In Figure 3, we visualize the application of the VML algorithm to an example fMRI time-series realization of the model described in Section 2.1 with true, but unknown, parameter values *β* = (2, −1)^*T*^ and *λ* = (−0.5, −2)^*T*^. As above, we used an imprecise prior for *β* by setting

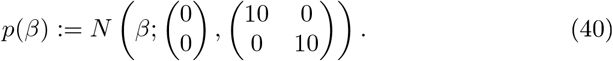

and set the initial covariance component estimate to *λ*^(1)^ = (0, 0)^*T*^. Panel A of Figure 3 depicts the prior distribution over *β* and the true, but unknown, value of *β*. Panel B depicts the variational distribution over *β* after convergence with a VML free energy convergence criterion of *δ* = 10^−3^. As in the VB scenario, given the imprecise prior distribution, this variational distribution falls close to the true, but unknown, value and convergence is usually achieved within 4 to 6 iterations. Panels C and D depict the VML free energy surface as a function of the variational parameter *m_β_* and the parameter *λ*, respectively. For the chosen prior distributions, the VML free energy surfaces displays a clear global maximum as a function of *m_β_*, while the maximum location as a function of *m_λ_* is located on an elongated crest.

**Figure 3:**
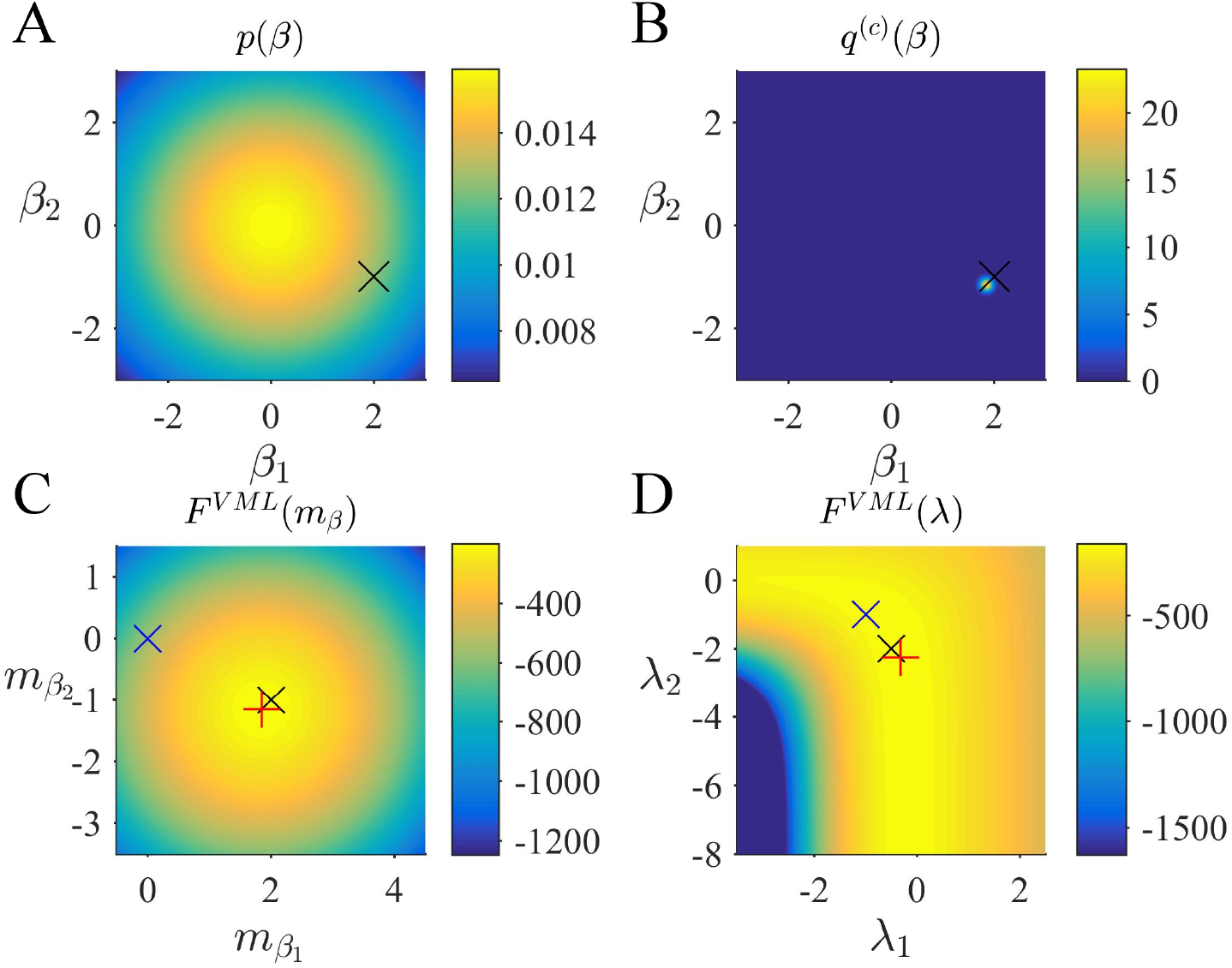
VML estimation. (A) Prior distribution *p*(*β*) with expectation *μ_β_* := (0, 0)^*T*^ and covariance Σ_*β*_ := 10*I*_2_. Here, and in all subpanels, the black x marks the true, but unknown, parameter value. (B) Variational approximation *q*^(*c*)^ (*β*) to the posterior distribution upon convergence of the algorithm. (C) VML free energy dependence on *m_β_*. The blue x indicates the prior expectation parameter and the red + marks the approximated posterior expectation parameter. (D) VML free energy dependence on *λ*. The blue x indicates the parameter value at algorithm initialization and the red + marks the parameter value upon algorithm convergence. For implementational details, please see *vbg_1.m*.

### 2.5 Restricted Maximum Likelihood (ReML)

ReML is commonly viewed as a generalization of the maximum likelihood approach, which in the case of the GLM yields unbiased, rather than biased, covariance component parameter estimates (Harville, 1977; Searle et al., 2009; Phillips et al., 2002). In this context and using our denotations, the ReML estimate *λ_ReML_* is defined as the maximizer of the ReML objective function

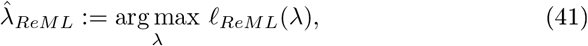

where

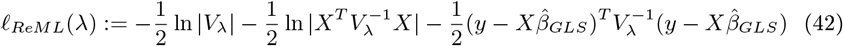

denotes the ReML objective function and

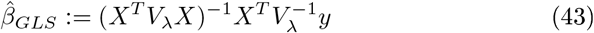

denotes the generalized least-squares estimator for *β*. Because 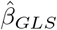 depends on *λ* in terms of *V_λ_*, maximizing the ReML objective function necessitates iterative numerical schemes. Traditional derivations of the ReML objective function, such as provided by LaMotte (2007) and Hocking (2013), are based on mixed-effects linear models and the introduction of a contrast matrix A with the property that *A^T^X* = 0 and then consider the likelihood of *A^T^y* after cancelling out the deterministic part of the model. In Supplementary Material S1.4 we show that, up to an additive constant, the ReML objective function also corresponds to the VML free energy under the assumption of an improper constant prior distribution for *β*, and an exact update of the VML free energy with respect to the variational distribution of *β*, i.e., setting *q*(*β*) = *pλ*(*β*|*y*). In other words, for the probabilistic model

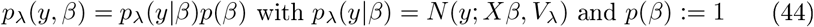

it holds that

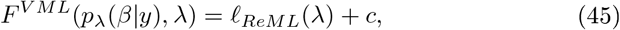

where

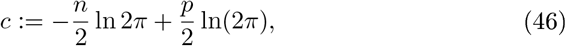

and thus

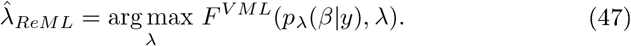

ReML estimation of covariance components in the context of the general linear model can thus be understood as the special case of VB, in which *β* is endowed with an improper constant prior distribution, the posterior distribution over *λ* is taken to be the Dirac measure *δ*_*λ*\*_, and the point estimate of *λ*\* maximizes the ensuing VML free energy under exact inference of the posterior distribution of *β*. In this view, the additional term of the ReML objective function with respect to the ML objective function obtains an intuitive meaning: 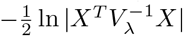 corresponds to the entropy of the posterior distribution *p_λ_*(*β*|*y*) which is maximized by the ReML estimate *λ_ReML_*. The ReML objective function thus accounts for the uncertainty that stems from estimating of the parameter *β* by assuming that is as large as possible under the constraints of the data observed.

In line with the discussion of VB and VML, we may define a ReML free energy, by which we understand the VML free energy function evaluated at 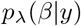 for the probabilistic model (44). In Supplementary Material S1.4, we show that this ReML free energy can be written as

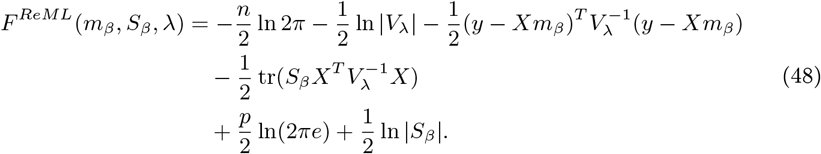

Note that the equivalence of eq. (48) to the constant-augmented ReML objective function of eq. (45) derives from the fact that under the infinitely imprecise prior distribution for *β* the variational expectation and covariance parameters evaluate to

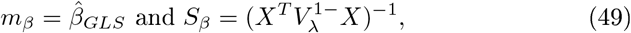

respectively. With respect to the general VML free energy, the ReML free energy is characterized by the absence of a term that penalizes the deviation of the variational parameter *m_β_* from its prior expectation, because the infinitely imprecise prior distribution *p*(*β*) provides no constraints on the estimate of *β*. To maximize the ReML free energy, we again derived a set of update equations which we document in Algorithm 3.

**Algorithm 3.**
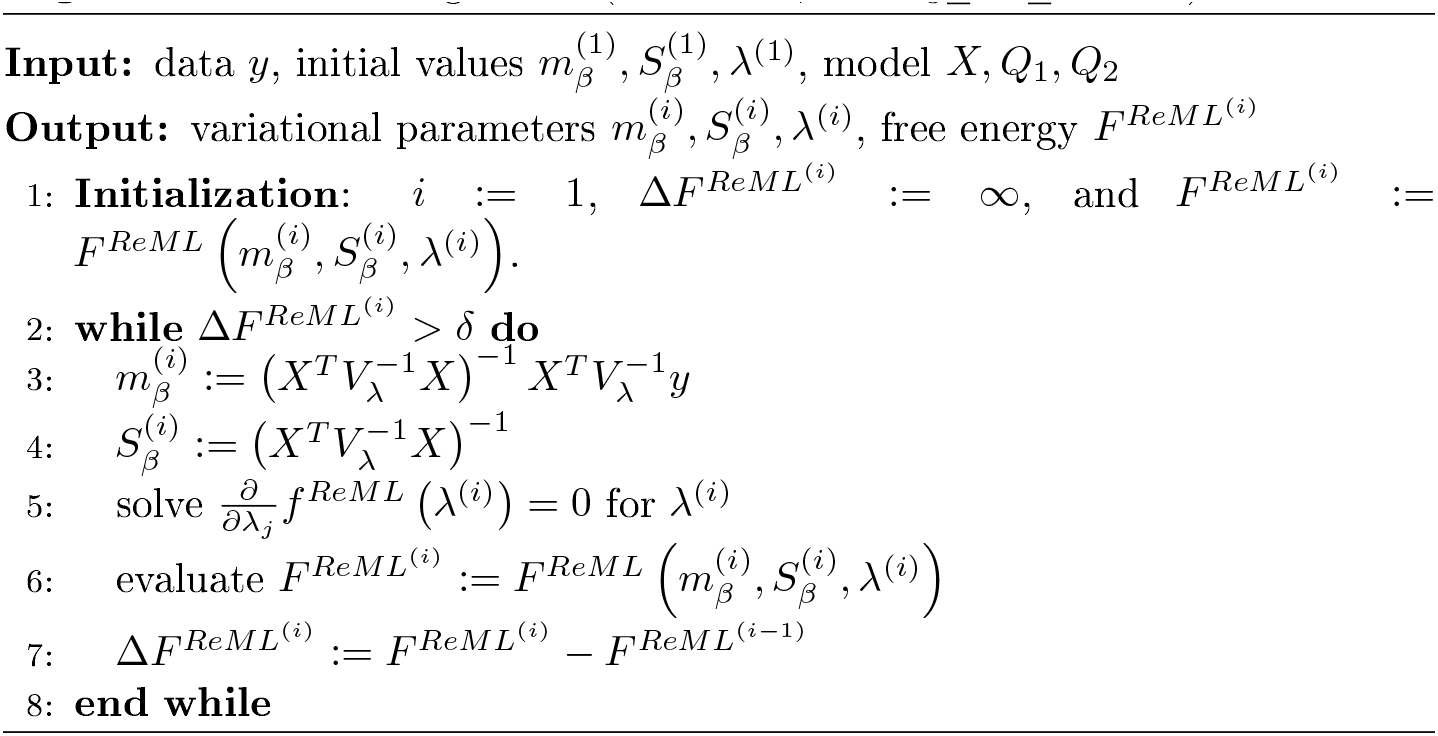
ReML Algorithm (for details, see *vbg_est_reml.m*)

In Figure 4, we visualize the application of the ReML algorithm to an example fMRI time-series realization of the model described in Section 2.1 with true, but unknown, parameter values *β* = (2, −1)^*T*^ and *λ* = (−0.5, −2)^*T*^. Here, we chose the *β* prior distribution parameters as the initial values for the variational parameters by setting

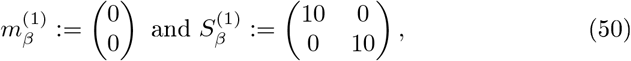

and as above, set the initial covariance component estimate to *λ*^(1)^ = (0, 0)^*T*^

**Figure 4:**
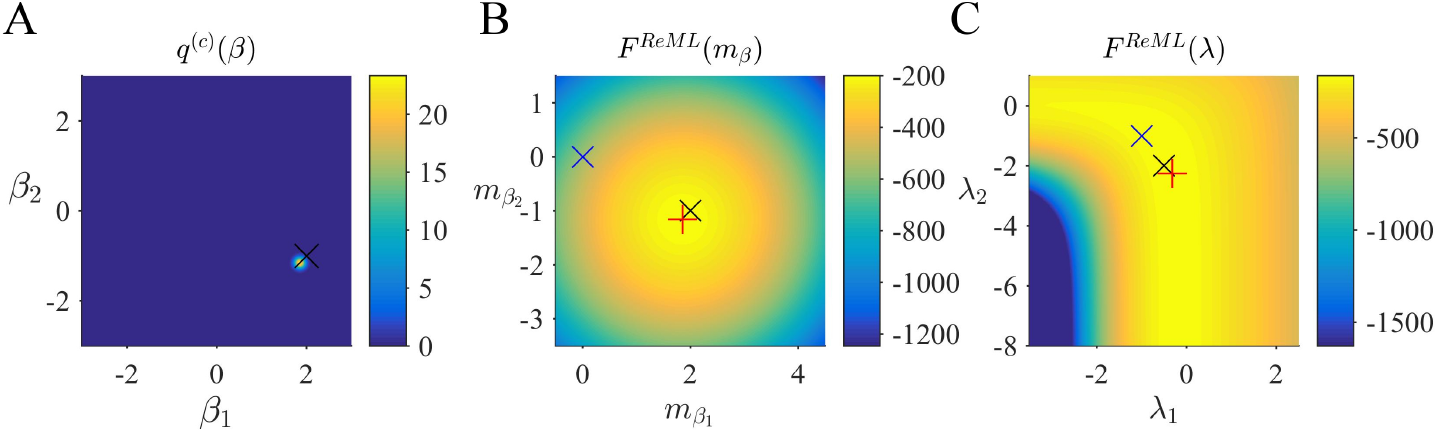
ReML estimation. (A) Variational distribution *q*^(*c*)^ (*β*) after convergence based on the initial values *m_β_* := (0, 0)^*T*^ and *S_β_* := 10*I*_2_ (convergence criterion *δ* = 10^−3^). Here, and in all subpanels, the black x marks the true, but unknown, parameter value. (B) ReML free energy dependence on *m_β_*. Here, and in Panel (C) the blue x indicates the parameter value at algorithm initialization and the red + marks the parameter value upon algorithm convergence. (C) ReML free energy dependence on *λ*. For implementational details, please see *vbg_1.m*.

Panel A of Figure 4 depicts the converged variational distribution over *β* and the true, but unknown, value of *β* for a ReML free energy convergence criterion of *δ* = 10^−3^. Panels B and C depict the ReML free energy surface as a function of the variational parameter *m_β_* and *λ*, respectively. Note that due to the imprecise prior distributions in the VB and VML scenarios, the resulting free energy surfaces are almost identical to the ReML free energy surface.

### 2.6 Maximum Likelihood (ML)

Finally, also the ML objective function can be viewed as the special case of the VB log marginal likelihood decomposition for variational distributions *q*(*β*) and *q*(*λ*) both conforming to Dirac measures. Specifically, as shown in Supplement Material S2 the ML estimate

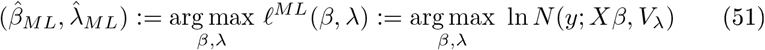

corresponds to the maximizer of the VML free energy for the probabilistic model

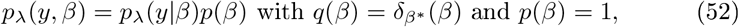

i.e. a Dirac measure *δ*_*β*\*_ for the variational distribution and an improper and constant prior density for the parameter *β*. Formally, we thus have

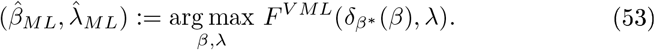

To align the discussion of ML with the discussion of VB, VML, and ReML, we may define the thus evaluated VML free energy as the *ML free energy*, which is just the standard log likelihood function of the GLM:

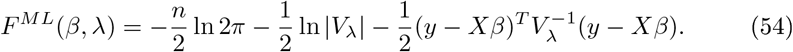

Note that the posterior approximation *q*(*β*) does not encode any uncertainty in this case, and thus the additional term corresponding to the entropy of this distribution in the ReML free energy vanishes for the case of ML. Finally, to maximize the ML free energy we again derived a set of update equations (Supplementary Material S1.5) which we document in Algorithm 4. In Figure 5, we visualize the application of this ML algorithm to an example fMRI time-series realization of the model described in Section 2.1 with true, but unknown, parameter values *β* = (2, −1)^*T*^ and *λ* = (−0.5, −2)^*T*^, initial parameter settings of *β*^(1)^ = (0, 0)^*T*^ and *λ*^(1)^ = (0, 0)^*T*^, and ML free energy convergence criterion *δ* = 10^−3^. Panel A depicts the ML free energy maximization with respect to *β*^(*i*)^ and Panel B depicts the ML free energy maximization with respect to *λ*^(*i*)^. Note the similarity to the equivalent free energy surfaces in the VB, VML, and ReML scenarios.

**Figure 5:**
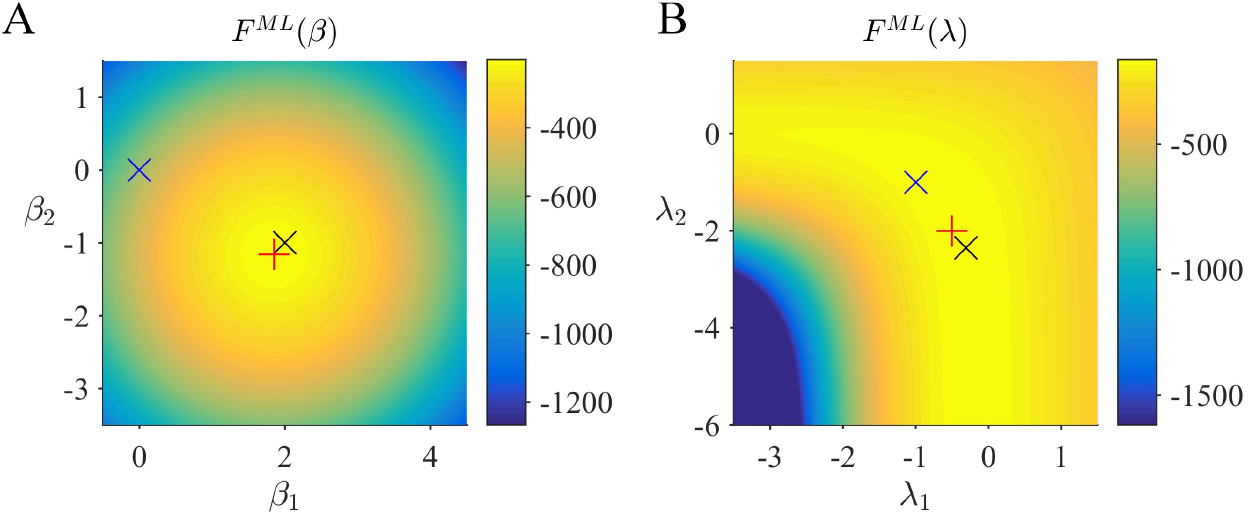
ML estimation. (A) ML free energy dependence on *β*. Here, and in Panel (B), the black x marks the true, but unknown parameter value, the blue x indicates the parameter value at algorithm initialization and the red + marks the parameter value upon algorithm convergence. (B) ML free energy dependence on *λ*. For implementational details, please see *vbg_1.m*.

In summary, in this section we have shown how VML, ReML, and ML estimation can be understood as special case of VB estimation. In the application to the GLM, the hierarchical nature of these estimation techniques yields a nested set of free energy objective functions, in which gradually terms that quantify uncertainty about parameter subsets are eliminated (cf. eqs. (28), (39), (48) and (54)). In turn, the iterative maximization of these objective functions yields a nested set of numerical algorithms, which assume gradually less complex formats (Algorithms 1 - 4). As shown by the numerical examples, under imprecise prior distributions, the resulting free energy surfaces and variational (expectation) parameter estimates are highly consistent across the estimation techniques. Finally, for all techniques, the relevant parameter estimates converge to the true, but unknown, parameter values after a few algorithm iterations.

**Algorithm 4.**
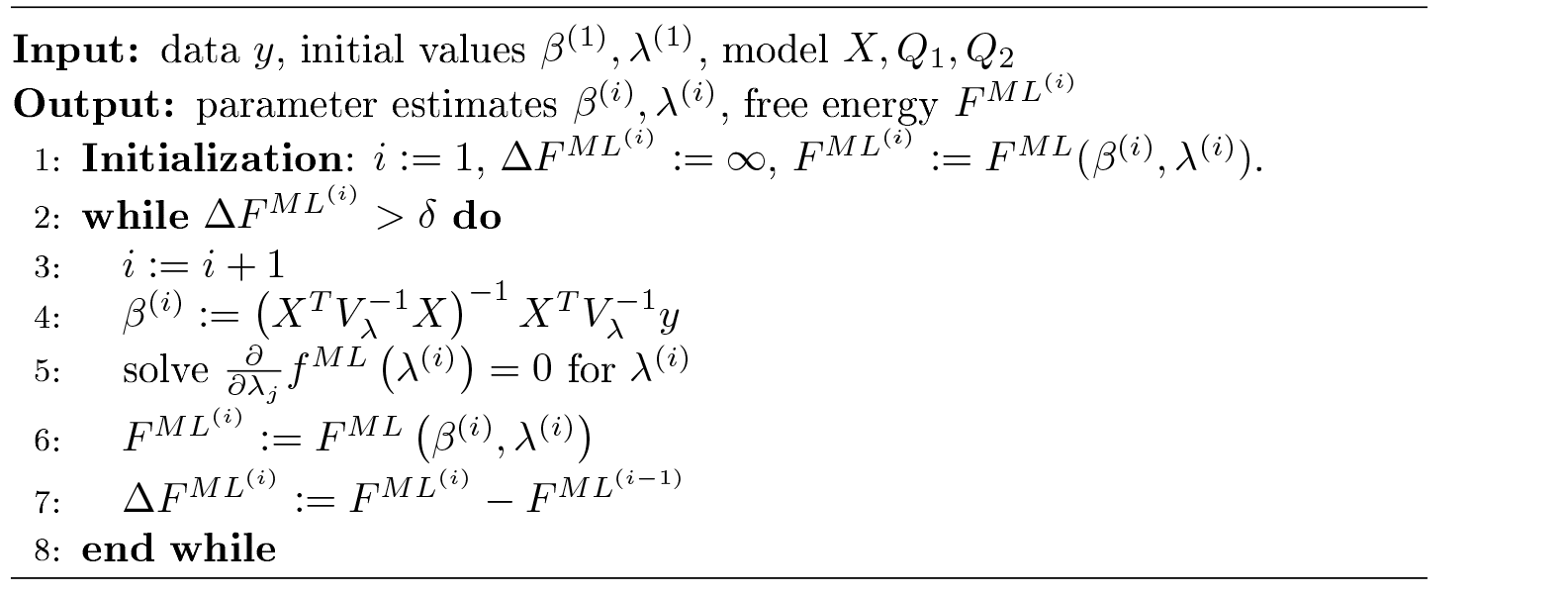
ML Algorithm (for details, see *vbg_est_ml.m*)

## 3 Applications

In Section 2 we have discussed the conceptual relationships and the algorithmic implementation of VB, VML, ReML, and ML in the context of the GLM and demonstrated their validity for a single simulated data realization. In the current section, we are concerned with their performance over a large number of simulated data realizations (Section 3.1) and their exemplary application to real experimental data (Section 3.2).

### 3.1 Simulations

Classical statistical theory has established a variety of criteria for the assessment of an estimator’s quality (e.g., Lehmann and Casella, 2006). Common, these criteria amount to the analytical evaluation of an estimators large sample behaviour. In the current section we adopt the spirit of this approach in simulations. To this end, we first capitalize on an objective Bayesian standpoint (Bernardo, 2003) by employing imprecise prior distributions to focus on the estimation techniques’ ability to recover the true, but unknown, parameters of the data generating model and the model structure itself. Specifically, we investigate the cumulative average and variance of the *β* and *λ* parameter estimates under VB, VML, ReML, and ML and the ability of each technique’s (marginal) likelihood approximation to distinguish between different data generating models. In a second step, we then demonstrate exemplarily how parameter prior specifications can induce divergences in the relative estimation qualities of the techniques.

#### Parameter Recovery

To study each estimation technique’s ability to recover true, but unknown, model parameters, we drew 100 realizations of the example model discussed in Section 2.1 and focussed our evaluation on the cumulative averages and variances of the converged (variational) parameter estimates 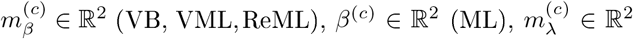, and 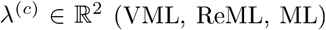. The simulations are visualized in Figure 6. Each panel column of Figure 6 depicts the results for one of the estimation techniques, and each panel row depicts the results for one of the four parameter values of interest. Each panel displays the cumulative average of the respective parameter estimate. Averages relating to estimates of *β* are depicted in blue, averages relating to estimates of *λ* are depicted in green. In addition to the cumulative average, each panel shows the cumulative variance of the parameter estimates as shaded area around the cumulative average line, and the true, but unknown, values *β* = (2, 1)^*T*^ and *λ* = (−0.5, −2)^*T*^ as grey line. Overall, parameter recovery as depicted here is within acceptable bounds and the estimates variances are tolerable. While there are no systematic differences in parameter recovery across the four estimation techniques, there are qualitative differences in the recovery of effect size and covariance component parameters. For all techniques, the recovery of the effect size parameters is unproblematic and highly reliable. The recovery of covariance component recovery, however, fails in a significant amount of approximately 15 - 20% of data realizations. In the panels relating to estimates of *λ* in Figure 6, these cases have been removed using an automated outlier detection approach (Grubbs, 1969). In the outlying cases, the algorithms converged to vastly different values, often deviating from the true, but unknown, values by an order of magnitude (for a summary of the results without outlier removal, please refer to Supplementary Material S3). To assess whether this behaviour was specific to our implementation of the algorithms, we also evaluated the defacto neuroimaging community standard for covariance component estimation, the *spm_reml.m* and *spm_reml_sc.m* functions of the SPM12 suite in the same model scenario. We report these simulations as Supplementary Material S4. In brief, we found a similar covariance component (mis)estimation behaviour as in our implementation.

**Figure 6:**
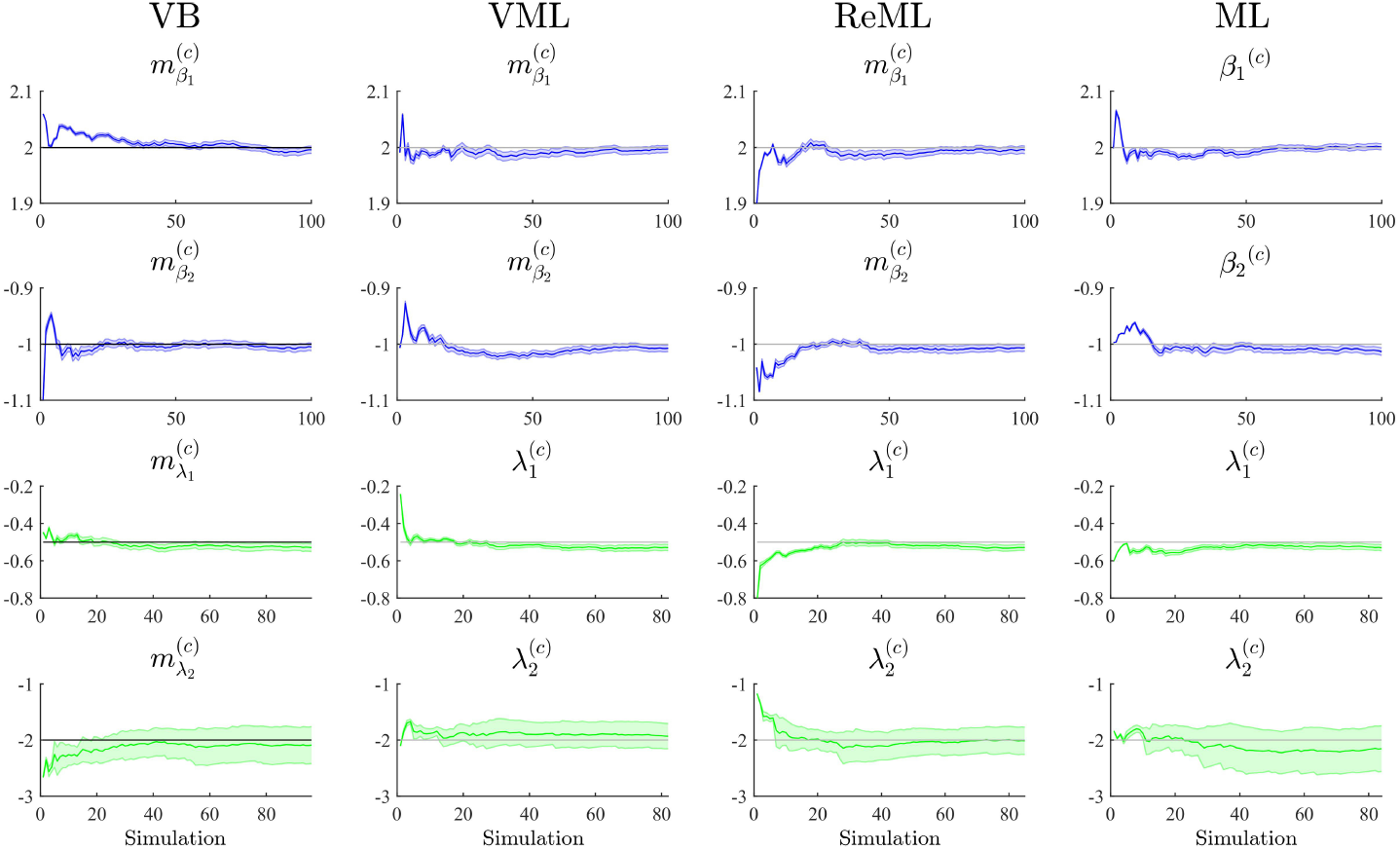
Parameter recovery. The panels along the figure’s columns depict the cumulative averages (blue/green lines), cumulative variances (blue/green shaded areas), and true, but unknown, parameter values (grey lines) for VB, VML, ReML, and ML estimation. Parameter estimates relating to the effect sizes *β* are visualized in blue, parameter estimates relating to the covariance components *λ* are visualized in green. The panels along the figure’s rows depict the parameter recovery performance for the subcomponents of the effect size parameters (row 1 and 2) and covariance component parameters (row 3 and 4), respectively. The covariance component parameter estimates are corrected for outliers as discussed in the main text. For implementational details, please see *vbg_2.m*.

Further research revealed that the relative unreliability of algorithmic covariance component estimation is a well-known phenomenon in the statistical literature (e.g., Groeneveld and Kovac, 1990; Boichard et al., 1992; Groeneveld, 1994; Foulley and van Dyk, 2000). We see at least two possible explanations in the current case. Firstly, we did not systematically explore the behaviour of the algorithmic implementation for different initial values. It is likely, that the number of estimation outliers can be reduced by optimizing, for each data realization, the algorithm’s starting conditions. However, also in this case, an automated outlier detection approach would be necessary to optimize the respective initial values. Secondly, we noticed already in the demonstrative examples in Section 2, that the free energy surface with respect to the covariance components is not as well-behaved as for the effect sizes. Specifically, the maximum is located on an elongated crest of the function, which is relatively at (see e.g. panel B of Figure 5) and hence impedes the straight-forward identification of the maximizing parameter value (see also Figure 4 of (Groeneveld and Kovac, 1990) for a very similar covariance component estimation objective function surface). In the Discussion section, we suggest a number of potential remedies for the observed outlier proneness of the covariance component estimation aspect of the VB, VML, ReML, and ML estimation techniques.

#### Model Recovery

Having established overall reasonable parameter recovery properties for our implementation of the VB, VML, ReML, and ML estimation techniques, we next investigated the ability of the respective techniques’ (marginal) log likelihood approximations to recover true, but unknown, model structures. We here focussed on the comparison of two data generating models that differ in the design matrix structure and have identical error covariance structures. Model MG1 corresponds to the first column of the example design matrix of Figure 1 and thus is parameterized by a single effect size parameter. Model MG2 corresponds to the model used in all previous applications comprising two design matrix columns. To assess the model recovery properties of the different estimation techniques, we generated 100 data realizations based on each of these two models with true, but unknown, effect size parameter values of *β*_1_ = 2 (MG1 and MG2) and *β*_2_ = −1 (MG2 only), and covariance component parameters *λ* = (−0.5, −2)^*T*^ (MG1 and MG2), as in the previous simulations. We then analysed each model’s data realizations with data analysis models that corresponded to only the single data-generating design matrix regressor (MA1) or both regressors (MA2) for each of the four estimation techniques.

The results of this simulation are visualized in Figure 7. For each estimation technique (panels), the average free energies, after exclusion of outlier estimates for the covariance component parameters, are visualized as bars. The data-generating models MG1 and MG2 are grouped on the x-axis and the data-analysis models are grouped by bar color (MA1 green, MA2 yellow). As evident from Figure 7, the correct analysis model obtained the higher free energy, i.e. log model evidence approximation, for both data-generating models across all estimation techniques. This difference was more pronounced when analysing data generated by model MG2 than when analysing data generated by model MG1. In this case, the observed data pattern is clearly better described by MA2. In the case of the data-generating model MG1, data analysis model MA2 can naturally account for the observed data by estimating the second effect size parameter to be approximately zero. Nevertheless, this additional model exibility is penalized correctly by all algorithms, such that the more parsimonious data analysis model MAI assumes the higher log model evidence approximation also in this case. We can thus conclude that model recovery is achieved satisfactorily by all estimation techniques. A more detailed decomposition of the average free energies into the respective free energy’s sum terms is provided in Supplementary Material S5.

**Figure 7:**
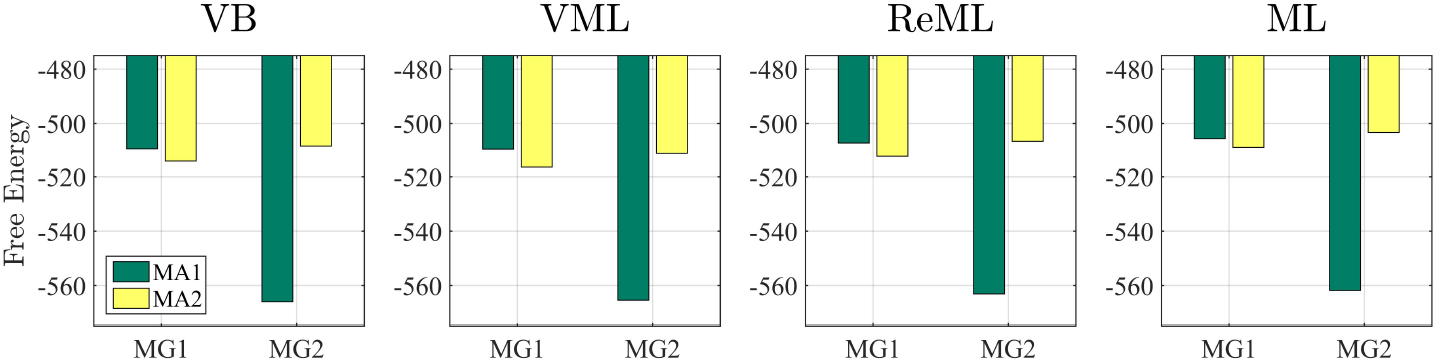
Model recovery. Each panel depicts the average free energies of the indicated estimation technique over 100 data realizations. Two data generating models (MG1 and MG2, panel x-axis) were used and analysed in a cross-over design with two data analysis models (MAI and MA2, bar color). MG1 and MAI comprise the same single column design matrix, and MG2 and MA2 comprise the same two column design matrix. Models MG1 and MAI are nested in MG2 and MA2. Across all estimation techniques, the correct data generating model is identified as indexed by the respective higher free energy log model evidence approximation. For implementational details, please see *vbg_3.m*.

#### Estimation quality divergences

Thus far, we have concentrated on the nested character of VML, ReML, and ML in VB and demonstrated that for the current model application the maximum-a-posteriori (MAP) estimates of VB and VML and the point estimates of ReML and ML are able to recover true, but unknown, parameter values. Naturally, the four estimation techniques differ in the information they provide upon estimation: VB estimates quantify posterior uncertainty about both effect size and covariance component parameters, VML estimates quantify posterior uncertainty about effect size parameters only, and ReML and ML do not quantify posterior uncertainty about either parameter class. Beyond these conceptual divergences, an interesting question concerns the qualitative and quantitative differences in estimation that result from the estimation techniques’ specific characteristics. In general, while the properties of ML estimates are fairly well understood from a classical frequentist perspective, the same cannot be said for the other techniques (e.g. Blei et al., 2016). We return to this point in the Discussion section. In the current section, we demonstrate divergences in the quality of parameter estimation that emerge in high noise scenarios, which are able to uncover prior distribution induced regularization effects. We demonstrate this for both effect size (Figure 8A) and covariance component parameters (Figure 8B) in the example model described in Section 2.1.

**Figure 8:**
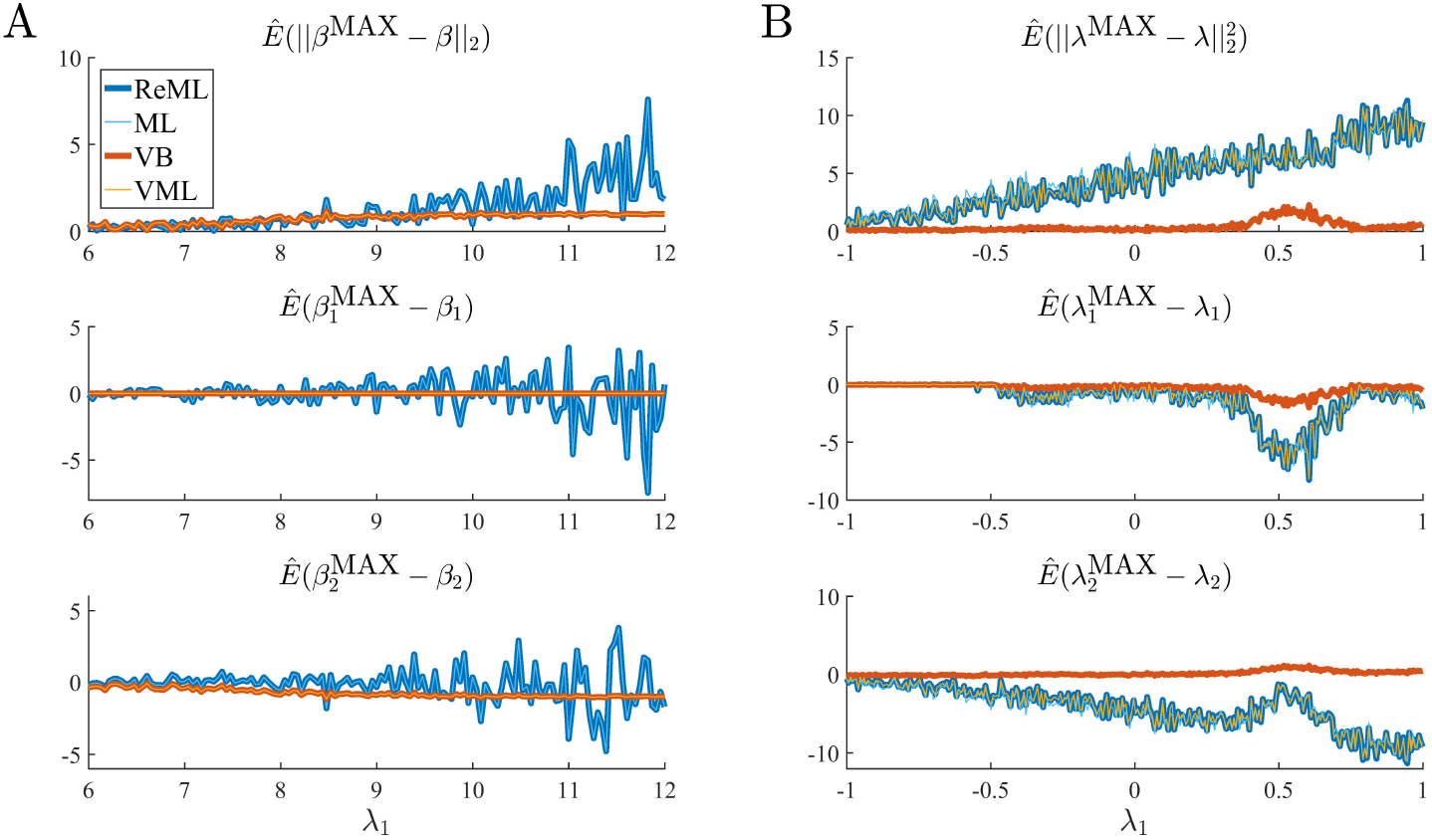
Estimation quality divergences. Each panel depicts the estimated RMSE and estimation bias for all four estimation techniques over a range of noise levels parameterized by *λ*_1_. The estimation techniques are color and linewidth coded. Panel A visualizes a simulation with focus on the effect size parameter estimates *β*, Panel B visualizes a simulation with focus on the covariance component parameters *λ* implementational details, please see *vbg_4.m*. Note that for Panel A, the results of VB and VML and the results of ReML and ML coincide, and for Panel B the results of ReML and VML coincide.

The panels in Figure 8A depict simulation estimates of the the root-mean-square-error (RMSE) 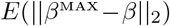 (uppermost panel) and biases of the effect size parameter entries 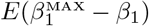 and 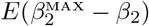 (middle and lowermost panel, respectively) over a range of values of the first covariance component parameter *λ*_1_. Here, 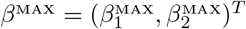 denotes the MAP estimates resulting under the VB and VML techniques, and the maximum (restricted) likelihood estimates resulting under ReML and ML, *β* denotes the true, but unknown, effect size parameter, *E*(·) denotes the expectation parameter, 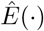 the estimation of an expectation by means of an average, and || · ||2 denotes the Euclidean norm of a vector. The results for the different estimation techniques are color-and linewidth-coded and were obtained under the following simulation: the true, but unknown, effect size parameter values were set to *β* = (1, 1)^*T*^ and the true, but unknown, parameter value of the second covariance component parameter was constant at *λ*_2_ = −2. Varying the true, but unknown, value *λ*_1_ on the interval [6, 12] thus increased the contribution of independent and identically distributed noise to the data. For each estimation technique, the respective effect size estimates were initialized as specified in Table 1. In brief, the estimates for *β*_1_ were initialized to the true, but unknown, value and *β*_2_ to zero. Crucially, VB and VML allow for the specification of prior distributions over *β*. Here, we used a precise prior covariance of Σ_*β*1_ = 10^−2^ and an imprecise variance of Σ_*β*2_ = 10^1^. Note that these algorithm parameters do not exist in ReML and ML. For each setting of *λ*_1_, 100 realizations of the model were obtained, subjected to all four estimation techniques, and the RMSE and biases estimated by averaging over realizations. The following pattern of results emerges: in terms of the RMSE (upper panel), VB and VML exhibit a more stable estimation of *β*, with a lower deviation from zero compared to the trend of ReML and ML estimates, at higher noise levels. In more detail, this pattern results from the following effects on the individual *β*_1_ and *β*_2_ estimates: first, for VB and VML, the estimates *β*_1_ exhibit virtually no biases, because their precise prior distribution fixes them at the true, but unknown value, (middle panel). For *β*_2_ this regularization of *β*_1_ results in more stable estimates as compared to ReML and ML, but for higher levels of noise also results in a downward bias (lowermost panel). Taken together, this simulation demonstrates, how, in the case of prior knowledge about the effect size parameters, the endowment of their estimates with precise priors in VB and VML can stabilize the overall effect size estimation and yield better estimates compared to ReML and ML.

**Table 1:** Parameter initialization for the simulations reported in Figure 8A and 8B design.

|  | VB |  |  |  | VML |  |  | ReML |  | ML |  |
| --- | --- | --- | --- | --- | --- | --- | --- | --- | --- | --- | --- |
| | $m_{\beta}^{(1)}$ | $S_{\beta}^{(1)}$ | $m_{\lambda}^{(1)}$ | $S_{\lambda}^{(1)}$ | $m_{\beta}^{(1)}$ | $S_{\beta}^{(1)}$ | $\lambda^{(1)}$ | $\beta^{(1)}$ | $\lambda^{(1)}$ | $\beta^{(1)}$ | $\lambda^{(1)}$ |
| 8A | $\begin{pmatrix} 1 \\ 0 \end{pmatrix}$ | $\begin{pmatrix} 0.01 & 0 \\ 0 & 10 \end{pmatrix}$ | $\begin{pmatrix} -1 \\ -1 \end{pmatrix}$ | $\begin{pmatrix} 10 & 0 \\ 0 & 10 \end{pmatrix}$ | $\begin{pmatrix} 1 \\ 0 \end{pmatrix}$ | $\begin{pmatrix} 0.01 & 0 \\ 0 & 10 \end{pmatrix}$ | $\begin{pmatrix} -1 \\ -1 \end{pmatrix}$ | $\begin{pmatrix} 1 \\ 0 \end{pmatrix}$ | $\begin{pmatrix} -1 \\ -1 \end{pmatrix}$ | $\begin{pmatrix} 1 \\ 0 \end{pmatrix}$ | $\begin{pmatrix} -1 \\ -1 \end{pmatrix}$ |
| 8B | $\begin{pmatrix} 0 \\ 0 \end{pmatrix}$ | $\begin{pmatrix} 10 & 0 \\ 0 & 10 \end{pmatrix}$ | $\begin{pmatrix} -1 \\ -1 \end{pmatrix}$ | $\begin{pmatrix} 10 & 0 \\ 0 & 10 \end{pmatrix}$ | $\begin{pmatrix} 0 \\ 0 \end{pmatrix}$ | $\begin{pmatrix} 10 & 0 \\ 0 & 10 \end{pmatrix}$ | $\begin{pmatrix} -1 \\ -1 \end{pmatrix}$ | $\begin{pmatrix} 0 \\ 0 \end{pmatrix}$ | $\begin{pmatrix} -1 \\ -1 \end{pmatrix}$ | $\begin{pmatrix} 0 \\ 0 \end{pmatrix}$ | $\begin{pmatrix} -1 \\ -1 \end{pmatrix}$ |

In a second simulation, visualized in Figure 8B, we investigated the interaction between prior regularization and estimation quality for the covariance component parameters. As in Figure 8A, the uppermost panel depicts the estimated RMSE for the *λ* parameters, and the middle and lowermost panels the biases of each component parameter. As in the previous simulation, the true, but unknown, effect size parameter values were set to *β* = (1, 1) and *λ*_2_ = −2 and *λ*_1_ was varied on the interval [−1, 1]. The initial parameters for each estimation technique are documented in Table 1. In brief, all effect size parameter estimates (expectations) were initialized to zero, and isotropic, imprecise prior covariance matrices were employed for VB and VML. The only estimation technique that endows *λ* estimates with a prior distribution is VB. Here, we employ the imprecise prior covariance Σ_*λ*_ := 10^1^*I*_2_, which is, however, precise enough to exert some stabilization effects: as shown in the uppermost panel of Figure 8B, only the RMSE of the VB technique remains largely constant over the investigated space of *λ*_1_ values, while for all other estimation techniques the RMSE increases linearly. Two things are noteworthy here. First, at the level of the *β* estimates all techniques perform equally well in a bias-free manner (data not shown). Second, the *λ*_1_ parameter space investigated includes a region (around 0.5) for which also the VB estimation quality declines, but recovers thereafter, suggesting an interaction between the structural model properties and the parameter regime. For the individual entries of *λ*, the decline in estimation quality in VML, ReML, and ML is not uniform: for *λ*_1_, the estimation quality remains largely constant up to the critical region around 0.5, whereas the estimation quality of *λ*_2_ deteriorates with increasing values of *λ*_1_ and recovers briefly in the critical region around 0.5. Note that for both simulations of Figure 8 we did not attempt to remove potential estimation outliers, because their definition in high noise scenarios is virtually impossible. It is thus likely, that the convergence failures observed in the first set of simulations contribute to the observed estimation errors. However, because these failures also afflict the VB and VML techniques which displayed improved estimation behaviour in the simulations reported in Figure 8, it is likely that the observed pattern of results is indicative of qualitative estimation divergences.

In summary, in the reported simulations we tried to evaluate our implementation of VB, VML, ReML, and ML estimation techniques for a typical nouroimaging data analysis example. In our first simulation set, we observed generally satisfactory parameter recovery for imprecise priors, with the exception of covariance component parameter recovery on a subset of data realizations. In our second simulation, we additionally observed satisfactory model recovery. In our last set of simulations, we probed for estimation quality divergences between the techniques and could show how regularizing prior distributions of the advanced estimation techniques VB and VML can aid to stabilize effect size and covariance component parameter estimation. Naturally, the reported simulations are conditional on our chosen model structure, the true, but unknown, parameter values, and the algorithm initial conditions (prior distributions), and hence not easily generalizable.

### 3.2 Application to real data

Having validated the VB, VML, ReML, and ML implementation in simulations, we were interested in their application to real experimental data with the main aim of demonstrating the possible parameter inferences that can (and cannot) be made with each technique. To this end, we applied VB, VML, ReML, and ML to a single participant fMRI data set acquired under visual checkerboard stimulation as originally reported in (Ostwald et al., 2010). In brief, the participant was presented with a single reversing left hemi-field checkerboard stimulus for 1 second every 16.5 to 21 seconds. These relatively long inter-stimulus intervals were motivated by the fact that the data was acquired as part of an EEG-fMRI study that investigated trial-by-trial correlations between EEG and fMRI evoked responses. Stimuli were presented at two contrast levels and there were 17 stimulus presentations per contrast level. 441 volumes of T2*-weighted functional data were acquired from 20 slices with 2.5 x 2.5 x 3 mm resolution and a TR of 1.5 seconds. The slices were oriented parallel to the AC-PC axis and positioned to cover the entire visual cortex. Data preprocessing using SPM5 included anatomical realignment to correct for motion artefacts, slice scan time correction, re-interpolation to 2 x 2 x 2 mm voxels, anatomical normalization, and spatial smoothing with a 5 mm FWHM Gaussian kernel. For full methodological details, please see (Ostwald et al., 2010).

To demonstrate the application of VB, VML, ReML, and ML to this data set, we used the SPM12 facilities to create a three-column design matrix for the mass-univariate analysis of voxel time-course data. This design matrix included HRF-convolved stimulus onset functions for both stimulus contrast levels and a constant offset. The design matrix is visualized in panel C of Figure 10. We then selected one slice of the preprocessed fMRI data (MNI plane z = 2) and used our implementation of the four estimation techniques to estimate the corresponding three effect size parameters 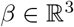 and the covariance component parameters 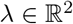 of the two covariance basis matrices introduced in Section 2.1 for each voxel. We focus our evaluation on the resulting variational parameter estimates of the effect size parameter *β*_1_, corresponding to the high stimulus contrast, and the first covariance component parameter *λ*_1_, corresponding to the isotropic error component. In line with the common practice in neuroimaging data analysis, no outlier removal was performed for the latter parameter. The results are visualized in Figures 9 and 10.

**Figure 9:**
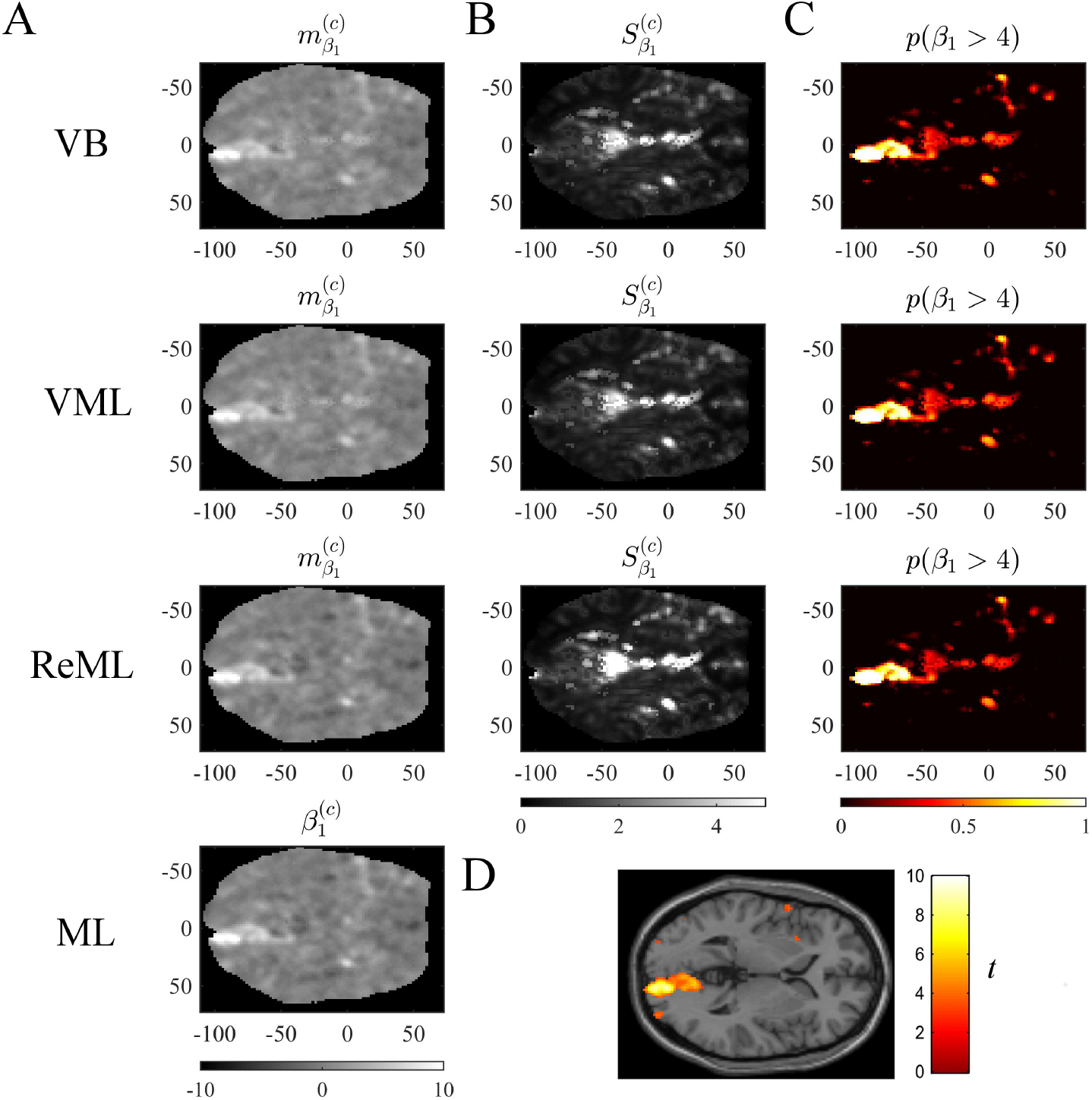
Effect size estimation. The figure panels depict the effect size parameter *β*_1_ estimation results of the VB, VML, ReML, and ML algorithm application to the analysis of a single-participant single-run fMRI data set. This effect size parameter captures the effect of high contrast left visual hemifield checkerboard stimuli as encoded by the first column of the design matrix shown in panel C of Figure 9. The first column (panel A) displays the converged expectation parameter estimates, the second column (panel B) the associated variance estimates, and the third column (C) the posterior probability for the true, but unknown, effect size parameter to assume values larger than 4. For visual comparison, panel D depicts the result of a standard GLM data analysis of the same data set using SPM12. For implementational details, please see *vbg_ 5.m*.

**Figure 10:**
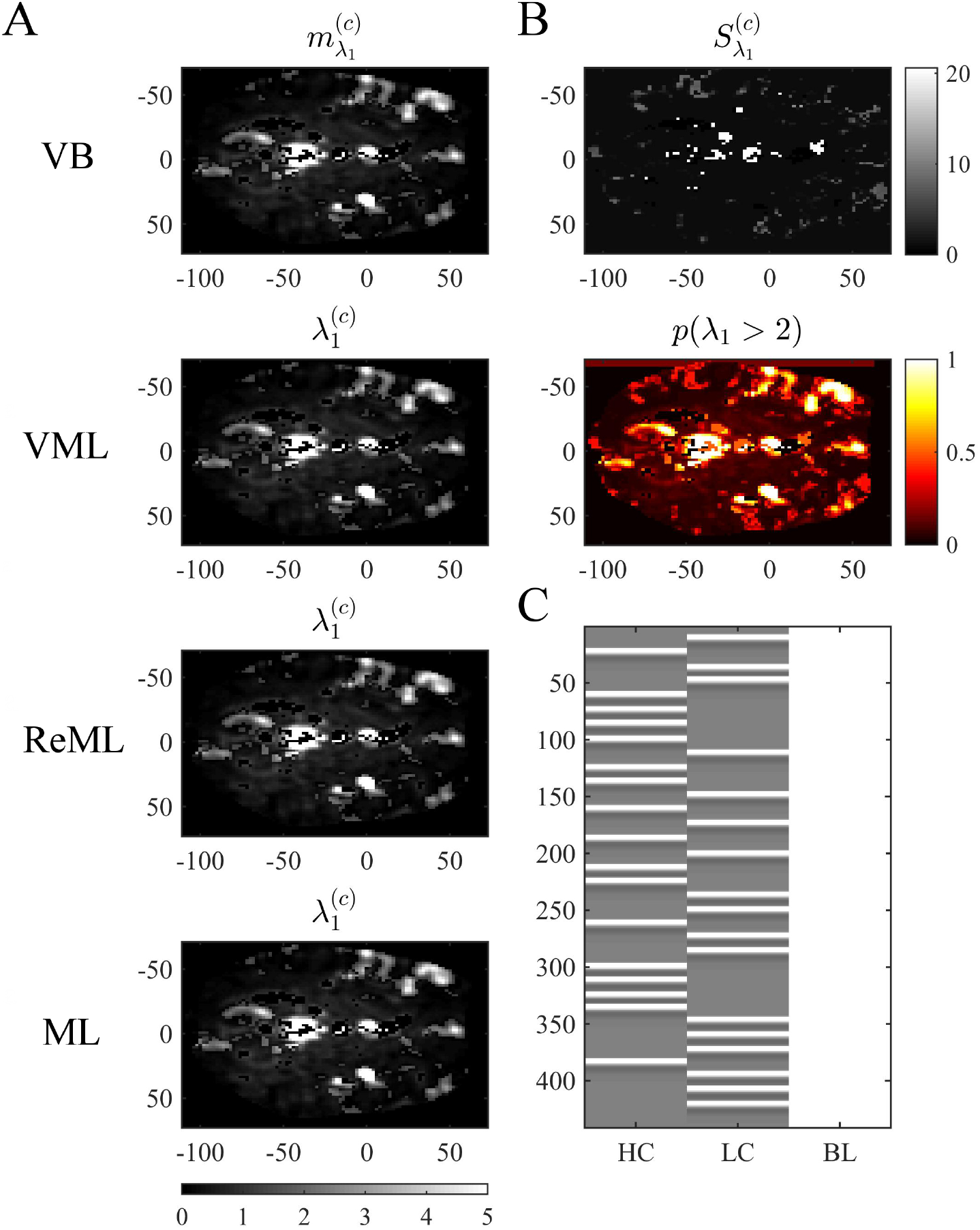
Covariance component parameter estimation. The figure panels depict the covariance component parameter *λ*_1_ estimation results of the VB, VML, ReML, and ML algorithm application to the analysis of a single-participant single-run fMRI data set. This covariance component parameter captures the effect of independently distributed errors. The first column (panel A) displays the converged (expectation) parameter estimates. The second column (panel B) displays the associated variance estimate and posterior probability for *λ*_1_ > 2, which is only quantifiable under the VB estimation technique. Panel C depicts the GLM design matrix that was used for the fMRI data analysis presented in Figures 8 and 9 (HC: high contrast stimuli regressor, LC: low contrast stimuli regressor, BL: baseline offset regressor). For implementational details, please see *vbg_5.m*.

Figure 9 visualizes the parameter estimates relating to the effect size parameter *β*_1_. The subpanels of Figure 10A depict the resulting two-dimensional map of converged variational parameter estimates, which differs only minimally between the four estimation techniques as indicated on the left of each panel. The variational parameter estimates are highest in the area of the right primary visual cortex, and lowest in the area of the cisterna ambiens/lower lateral ventricles. Panel B depicts the associated variational covariance parameter 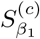, i.e., the first diagonal entry of the of the variational covariance matrix 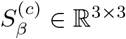. Here, the highest uncertainty is observed for ventricular locations and the right medial cerebral artery. Overall, the uncertainty estimates are marginally more pronounced for the VB and VML techniques compared to the ReML estimates. Note that the ML technique does not quantify the uncertainty of the GLM effect size parameters. Based on the variational parameters 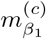 and 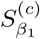, Panel C depicts the probability that the true, but unknown, effect size parameter is larger than *η* = 4, i.e.

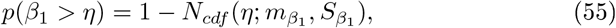

where *N_cdf_* denotes the univariate Gaussian cumulative density function. Here, the stimulus-contralateral right hemispheric primary visual cortex displays the highest values and the differences between VB, VML, and ReML are marginal. For comparison, we depict the result of a classical GLM analysis with contrast vector *c* = (1, 0, 0)^*T*^ at an uncorrected cluster-defining threshold of *p* < 0.001 and voxel number threshold of *k* = 0 overlaid on the canonical single participant T1 image in 9D. This analysis also identifies the right lateral primary visual cortex as area of strongest activation - but in contrast to the VB, VML, and ReML results does not provide a visual account of the uncertainty associated with the parameter estimates and ensuing T-statistics. In summary, the VB, VML, and ReML-based quantification of effect sizes and their associated uncertainty revealed biologically meaningful results.

Figure 10 visualizes the variational expectation parameters relating to the effect size parameter *λ*_1_. Here, the subpanels of Figure 10A visualize the variational (expectation) parameters across the four estimation techniques. High values for this covariance component are observed in the areas covering cerebrospinal fluid (cisterna ambiens, lateral and third ventricles), lateral frontal areas, and the big arteries and veins. Notably, also in right primary visual cortex, the covariance component estimate is relatively large, indicating that the design matrix does not capture all stimulus-induced variability. The only estimation technique that also quantifies the uncertainty about the covariance component parameters is VB. The results of this quantification are visualized in 10B. The first subpanel visualizes the variational covariance parameter 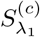, i.e., the first diagonal entry of the variational covariance matrix 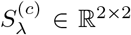. The second subpanel visualizes the probability that the true, but unknown, covariance component parameter *λ* is larger than *η* = 2, i.e.

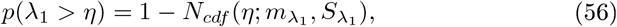

which, due to the relatively low uncertainty estimates S_λ1_ shows high similarity with the variational expectation parameter map. In summary, our exemplary application of VB, VML, ReML, and ML to real experimental data revealed biologically sensible results for both effect size and covariance component parameter estimates.

## 4 Discussion

In this technical study, we have reviewed the mathematical foundations of four major parametric statistical parameter estimation techniques that are routinely employed in the analysis of neuroimaging data. We have detailed, how VML (expectation-maximization), ReML, and ML parameter estimation can be viewed as special cases of the VB paradigm. We summarize these relationships and the non-technical application scenarios in which each technique corresponds to the method of choice in Figure 11. Further, we have provided a detailed documentation of the application of these four estimation techniques to the GLM with non-spherical, linearly decomposable error covariance, a fundamental modelling scenario in the analysis of fMRI data. Finally, we validated the ensuing iterative algorithms with respect to both simulated and real experimental fMRI data. In the following, we relate our exposition to previous treatments of similar topic matter, discuss potential future work on the qualitative properties of VB parameter estimation techniques, and finally comment on the general relevance of the current study.

**Figure 11:**
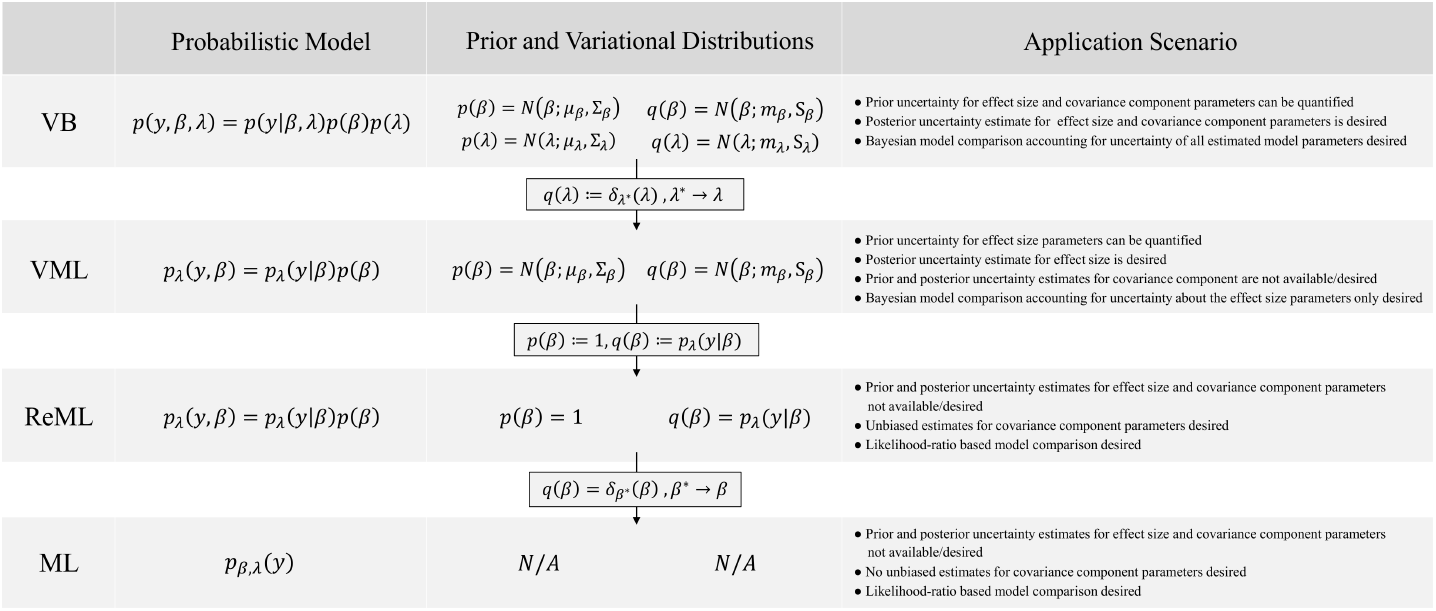
VB, VML, ReML, and ML relationships and application scenarios. N/A denotes non-applicable.

The relationships between VB, VML, ReML, and ML have been previously pointed out in Friston et al. (2002a) and Friston et al. (2007). In contrast to the current study, however, Friston et al. (2002a) and Friston et al. (2007) focus on high-level general results and provide virtually no derivations. Moreover, when introducing VB in Friston et al. (2007), the GLM with non-spherical, linearly decomposable error covariance is treated as one of a number of model applications and is not studied in detail across all estimation techniques. From this perspective, the current study can be understood as making many of the implicit results in Friston et al. (2002a) and Friston et al. (2007) explicit and filling in many of the detailed connections and consequences, which are implied by Friston et al. (2002a) and Friston et al. (2007). The relationship between VB and VML has been noted already from outset of the development of the VB paradigm (Beal, 2003; Beal and Ghamarani, 2003). In fact, VB was originally motivated as a generalization of the EM algorithm (Neal and Hinton, 1998; Attias, 2000). However, these treatments do not provide an explicit derivation of VML from VB based on the Dirac measure and do not make the connection to ReML. Furthermore, these studies do not focus on the GLM and its application in the analysis of fMRI data. Finally, a number of treatises have considered the application of VB to linear regression models (e.g., Bishop, 2006; Murphy, 2012; Tzikas et al., 2008). However, these works do not consider non-spherical linearly decomposable error covariance matrices and also do not make the connection to classical statistical estimation using ReML for functional neuroimaging. Taken together, the current study complements the existing literature with its emphasis on the mathematical traceability of the relationship between VB, VML, ReML, and ML, its focus on the GLM application, and its motivation from a functional neuroimaging background.

### Estimator quality

Model estimation techniques yield estimators. Estimators are functions of observed data that return estimates of true, but unknown, model parameters, be it the point-estimates of classical frequentist statistics or the posterior distributions of the Bayesian paradigm (e.g., Wasserman, 2010). An important issue in the development of estimation techniques is hence the quality of estimators to recover true, but unknown, model parameters and model structure. While this issue re-appears in the functional neuroimaging literature in various guises every couple of years (e.g., Vul et al., 2009a; Eklund et al., 2016a), often accompanied by some flurry in the field (e.g., Nichols and Poline, 2009; Vul et al., 2009b; Abbott, 2009; Eklund et al., 2016b; Miller, 2016), it is perhaps true to state that the systematic study of estimator properties for functional neuroimaging data models is not the most matured research field. From an analytical perspective, this is likely due to the relative complexity of functional neuroimaging data models as compared to the fundamental scenarios that are studied in mathematical statistics (e.g., Shao, 2003). In the current study, we used simulations to study both parameter and model recovery, and while obtaining overall satisfiable results, we found that the estimation of covariance component parameters can be deficient for a subset of data realizations. As pointed out in Section 3, this finding is not an unfamiliar result in the statistical literature (e.g., Groeneveld and Kovac, 1990; Boichard et al., 1992; Groeneveld, 1994; Harville, 1977). We see two potential avenues for improving on this issue in future research. Firstly, there exist a variety of covariance component estimation algorithm variants in the literature (e.g., Gilmour et al., 1995; Witkovsky, 1996; Thompson and Mäntysaari, 1999; Foulley and van Dyk, 2000; Misztal, 2008) and research could be devoted to applying insights from this literature in the neuroimaging context. Secondly, as the deficient estimation primarily concerns the covariance component parameter that scales the AR(1) + WN model covariance basis matrix, it remains to be seen, whether the inclusion of a variety of physiological regressors in the deterministic aspect of the GLM will eventually supersede the need for covariance component parameter estimation in the analysis of firstlevel fMRI data altogether (e.g., Glover et al., 2000; Lund et al., 2006). Finally, we presented the application of VB, VML, ReML, and ML in the context of fMRI time-series analysis. As pointed out in Section 1, the very same statistical estimation techniques are of eminent importance for a wide range of other functional neuroimaging data models. Moreover, together with the GLM, they also form a fundamental building block of model-based behavioural data analyses as recently proposed in the context of computational psychiatry (e.g., Montague et al., 2012; Stephan et al., 2016a,b,c; Schwartenbeck and Friston, 2016) and recent developments in the analysis of big data (e.g., Allenby et al., 2014; Ghahramani, 2015).

On a more general level, the relative merits of the parameter estimation techniques discussed herein form an important field for future research. Ideally, the statistical properties of estimators resulting from variational approaches were understood for the model of interest, and known properties of their specialized cases, such as the bias-free covariance component parameter estimation under ReML with respect to ML, would be deducible from these. However, as pointed out by Blei et al. (2016), the statistical properties of variational approaches are not yet well understood. Nevertheless, there exists a few results on the statistical properties of variational approaches, typically in terms of the variational expectations upon convergence and for fairly specific model classes. Of relevance for the model class considered herein is the recent work by You et al. (2014), who could show the consistency of the variational expectation in the frequentist sense, albeit for spherical covariance matrices and a gamma distribution for the covariance component parameter. For a broader model class with posterior support in real space (including the current model class of interest), Westling (2017) have worked towards establishing the consistency and asymptotic normality of variational expectation estimates. Finally, a number of authors have addressed consistency and asymptotic properties in selected model classes, such as Poisson-mixed effect models, stochastic block models, and Gaussian mixture models (Hall et al., 2011; Celisse et al., 2012; Bickel et al., 2013; Wang et al. 2006).

In summary, understanding the qualitative statistical properties of variational Bayesian estimators and their relative merits with respect to more specialized approaches forms a burgeoning field of research. New impetus in this direction may also arise from recent attempts to understand the properties of deep learning algorithms from a probabilistic variational perspective (Gal and Ghahramani, 2017).

### Conclusion

To conclude, we believe that the mathematization and validation of model estimation techniques employed in the neuroimaging field is an important endeavour as the field matures. With the current work, we attempted to provide a small step in this direction. We further hope to be able to contribute to a better understanding of the statistical properties of the parameter estimation techniques for neuroimaging-relevant model classes in our future work.

## Supplementary Material

Contents
**S1 Free energy algorithm derivations**

S1.1 Preliminaries
S1.2 Variational Bayes
S1.3 Variational maximum likelihood
S1.4 Restricted maximum likelihood
S1.5 Maximum likelihood
**S2 Foundations of variational Bayes**

S2.1 Preliminaries
S2.2 Entropies of distributions of random vectors
S2.3 Probability-theoretic variational Bayes
**S3 Cumulative averages without outlier removal**
**S4 SPM12 ReML estimation**
**S5 Model recovery free energy contributions**

## S1 Free energy algorithm derivations

In this section, we evaluate the VB, VML, ReML, and ML free energies for the GLM and derive update equations for their maximization. The notation follows the applied approach used in the main text. We commence with some remarks on additional notation and matrix differentation.

### S1.l Preliminaries

#### Expectations

To ease the notation, we will often write the expectation of a function *f* of random variable *x* under the probability distribution *p(x)* using the expectation operator

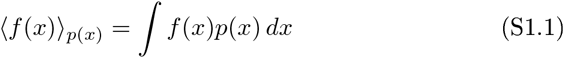

Furthermore, on numerous occasions, we require the following property of expectations of multivariate random variables 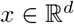 under normal distributions: for 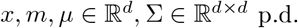 and 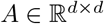 it holds that

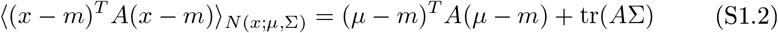

(see e.g. Petersen and Pedersen (2012), eq. (380)).

#### Gradient and Hessian

The gradient and Hessian of a real-valued function

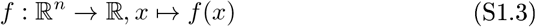

evaluated at a point 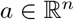 will be denoted by

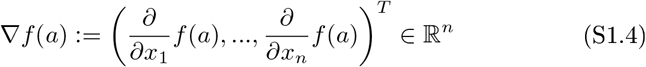

and

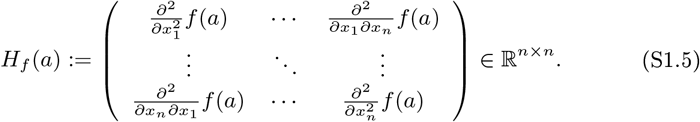

When it eases the notation, we also occasionally denote the partial derivative of *f* with respect to *x_i_* evaluated at 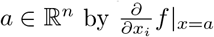.

#### Matrix differentiation

The following matrix differentiation rules are used in the subsequent derivations (Petersen and Pedersen, 2012). For a matrix *A* depending on a scalar parameter *x*, we have

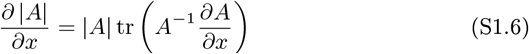

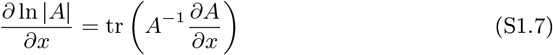

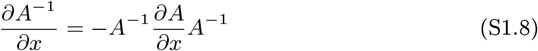

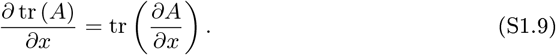

For a matrix *A* depending on a two-dimensional vector *x = (x_1_, x_2_)*, the second-order partial derivatives of its inverse are

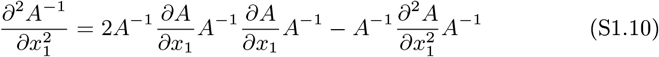

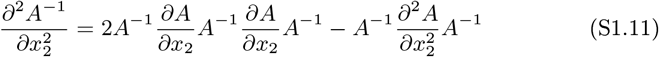

and

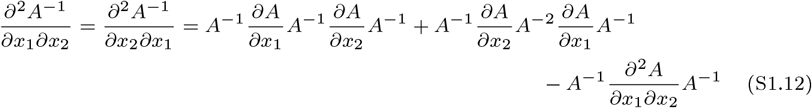

assuming that *A* has continuous second derivatives, such that the symmetry of second-order derivatives (Schwarz’s theorem) holds. For the update equations of the matrix parameters *S_β_* and *S_λ_*, we also need to compute derivatives regarding matrices. We have

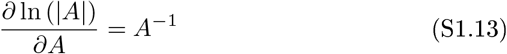

and for matrices *A* Mid *B* of matching dimensions

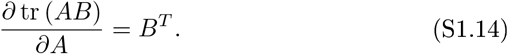

### S1.2 Variational Bayes

#### Evaluation of the VB free energy

To evaluate the VB free energy, we first rewrite it from its definition in eq. (20) in the main text as follows

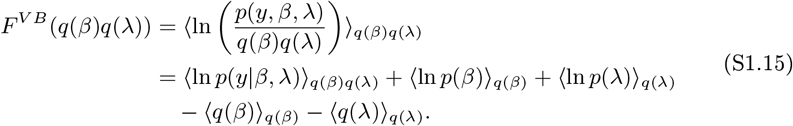

Using (S1.2), the second and third term on the right-hand side of (S1.15) can be evaluated exactly, yielding

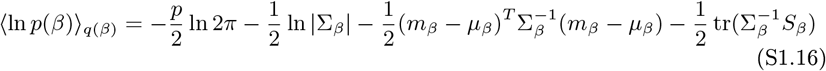

and

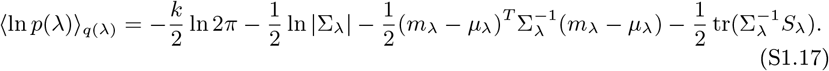

corresponding to terms 6 - 13 of eq. (28) in the main text. The fourth and the fifth term on the right-hand side of (S1.15) correspond to the entropies of the variational distributions, which given their Gaussian form are given as function of their respective covariance matrices (e.g., Bishop, 2006)

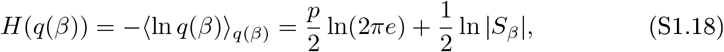

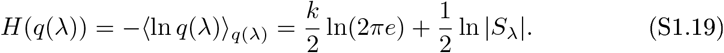

Eqs. (S1.18) and (S1.19) correspond to terms 14 to 16 of eq. (28) in the main text.

Finally, we consider the first term of (S1.15). Based on the definition of 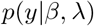, the expectation with respect to *q(β)* can be evaluated exactly, yielding

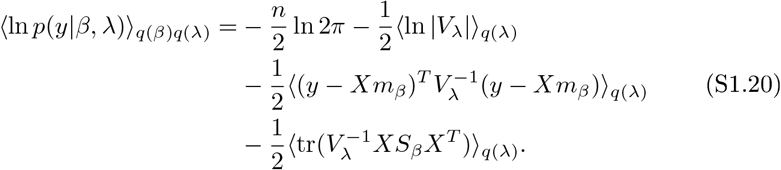

To make it possible to evaluate the remaining expectations, we use a second order Taylor approximation. Let

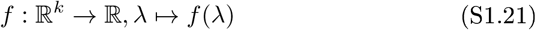

denote a real-valued function of λ. Then

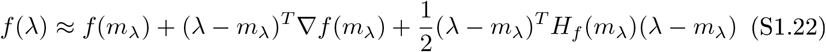

in the vicinity of *m*_λ_. If *q* (λ) is sufficiently narrow, that is, if most of its mass is concentrated close to *m*_λ_, we can thus approximate

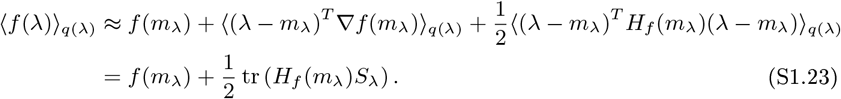

This approximation needs to be applied to all expectations in equation (S1.20). Thus, using the linearity of the trace to subsume all Hessian matrices into

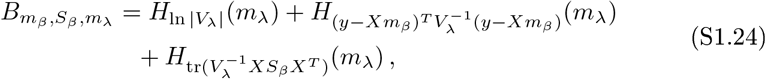

thereby pooling the second-order terms, we arrive at terms 1 - 5 of equation (28) in the main text, and the derivation is complete.

#### Evaluation of B_m_β_, S_β_, m_λ__

To estimate the VB free energy in practice, the Hessian matrices on the right-hand side of (S1.24) have to be evaluated. For the linear form of the error covariance matrix

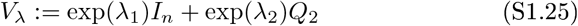

the three Hessian matrices of (S1.24) can be evaluated analytically:

• *H*_ln |*V*_λ_|_

Using (S1.7), the first order partial derivatives are given by

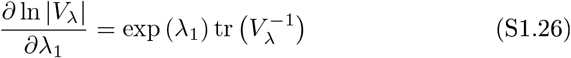

and

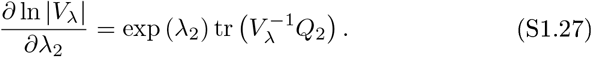

Exploiting the linearity of the trace operator (S1.9) and using (S1.8) for the derivative of the inverse yields the second order partial derivatives:

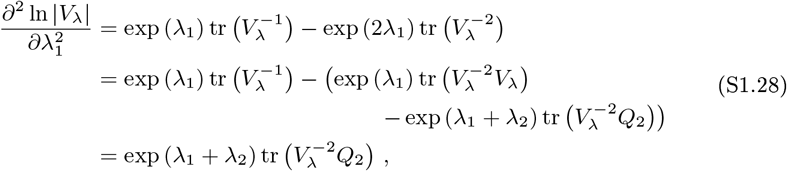

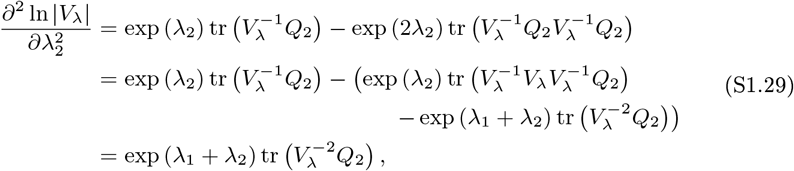

and

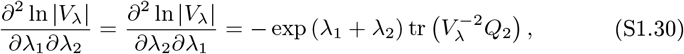

where in the last equation we used that the trace is invariant under cyclic permutations, e.g. tr (*ABC*) = tr (*CAB*) = tr (*BCA*).

• 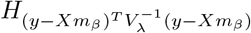

The Hessian matrix of 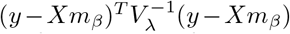 only depends on the second order partial derivatives of the inverse of *V*_λ_

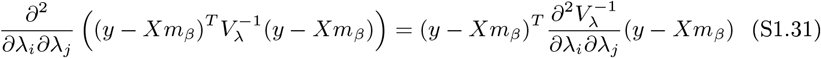

for *i, j* ∈ {1, 2}. Applying (S1.10) to (S1.12) yields

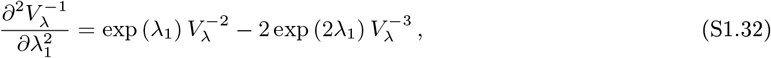

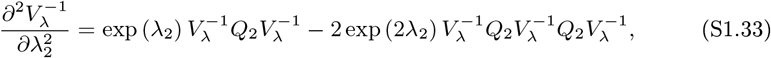

and

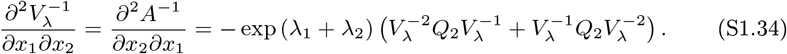

• 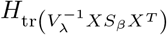

Due to the linearity of the trace Operator, we have

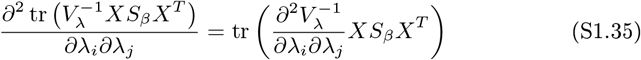

for *i, j* ∈ {1, 2}. Thus we only have to use (S1.32) to (S1.34).

Notably, the evaluation of these Hessian matrices will necessitate the inversion of *V*_λ_ on every iteration of the optimization algorithm. This inversion can be performed efficiently using the diagonalized form of *Q*_2_. As *Q*_2_ is a real, symmetric matrix by design, there exists a diagonalized form given by 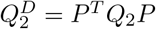, where *P* is a unitary transformation matrix (*P^T^ = P*^-1^). The entries *l_i_, i* ∈ {1,…, *n*} of 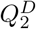 are the eigenvalues of *Q*_2_. We thus have

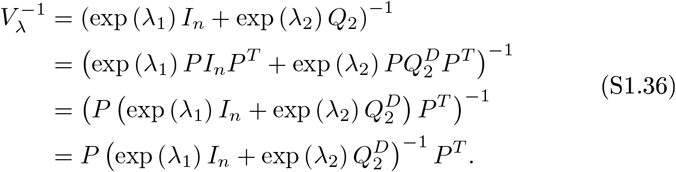

As exp 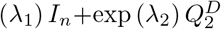 is a diagonal matrix, its inverse is easily evaluated, and the diagonalizing matrix *P* only needs to be computed once for any given *Q*_2_

#### The VB free energy update equations

In this section, we consider the iterative maximization of the VB free energy function with respect to its vector and matrix parameters *m_β_, S_β_, m_λ_* and *S*_λ_. In each case, we identify the relevant subpart of the VB free energy function depending on the respective parameter, evaluate its gradient with respect to the parameter in question, set the gradient to zero, and, if possible, solve the ensuing equation for a parameter update equation. To emphasize the iterative character of this endeavour, we use the superscript (*i*) to denote the values of parameters at a given algorithm iteration.

We consider the update with respect to *S*_λ_ first. The relevant subpart of 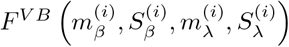 depending on *S_λ_* is given by

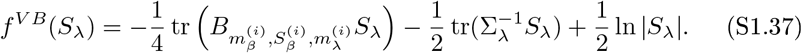

Using the identities (S1.13), (S1.14), and considering that 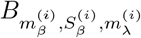 and 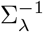 are symmetric, evaluation of the gradient of *f^VB^* results in

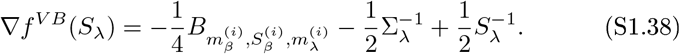

Setting the gradient to zero and solving for the parameter update 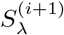 then yields

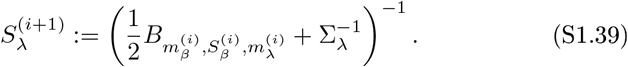

Note that with the linearity properties of the trace operator, this update equation implies as a result, that the sum of the two trace terms involving *S*_λ_ in the VB free energy (equation (28) of the main text) evaluates to 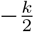 and the term 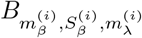 does not need to be considered when deriving the update equations for *m_β_, S_β_*, and *m_λ_*.

Next, the relevant subpart of 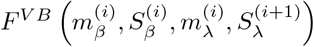 depending on *m_β_* is given by

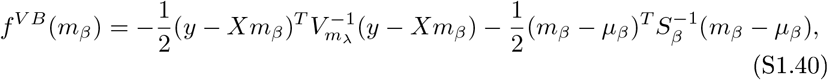

where we omitted iteration superscripts for visual clarity. With (S1.2), the gradient of *f^VB^(m_β_)* is given by

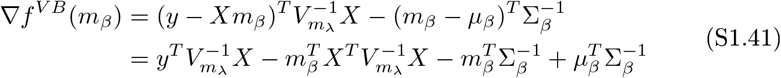

Setting the gradient to zero then yields the update equation

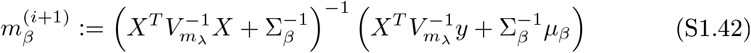

Analogously, the relevant subpart of 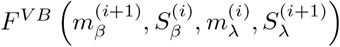 depending on *S_β_* is given by

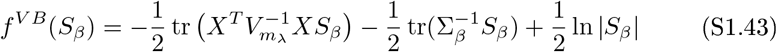

with gradient

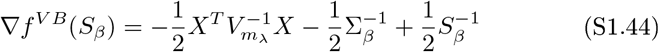

and the resulting update equation

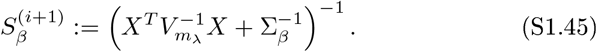

Note that the update equations (S1.42) and (S1.45) conform to the wellknown closed-form expressions for Bayesian inference in the conjugate Gaussian model (cf. eq. (9) of the main text), with the difference of the parametric dependence of the error covariance matrix on 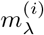.

Finally, the relevant subpart of 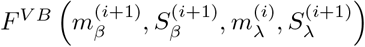 depending on *m*_λ_ is given by, again omitting iteration superscripts for visual clarity,

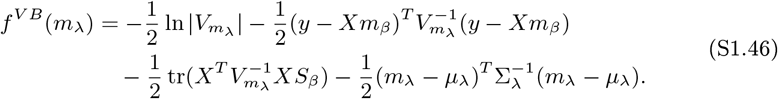

Evaluation of entries 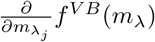 of the gradient 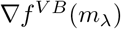 yields

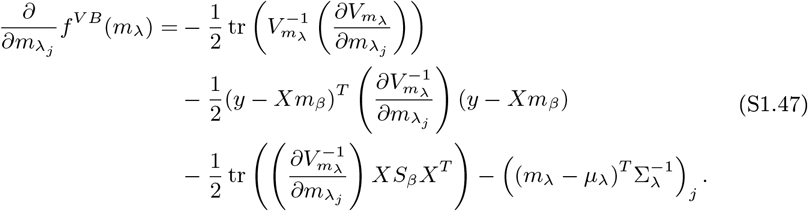

The evaluation of these entries for the two-component linear error covariance (S1.25) then yields

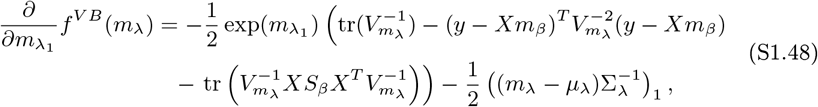

and

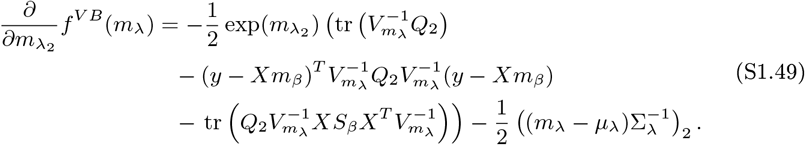

Lastly, to determine the value 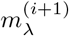 for which

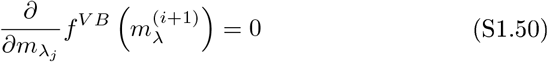

for *j* = 1, 2, we employ the routine *fsolve.m* provided by Matlab (MATLAB and Optimization Toolbox Release 2014b, The MathWorks, Inc., Natick, Massachusetts, United States). This function implements a trust-region dogleg algorithm for the minimization of nonlinear real-valued functions of multiple variables (Coleman and Li, 1996; Nocedal and Wright, 2006).

### S1.3 Variational maximum likelihood

#### Evaluation of the VML free energy

The VML free energy is defined as

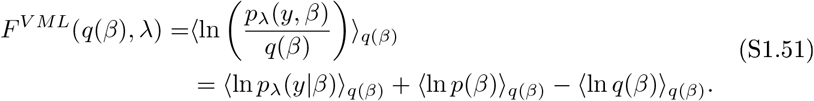

The latter two terms on the right-hand side of (S1.51) have been evaluated in Section S1.2. The first term can be evaluated using (S1.2), yield

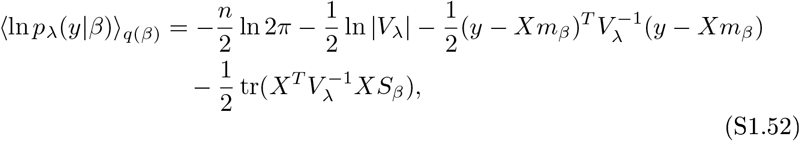

which completes the derivation of the VML free energy as eq. (39) of the main text.

#### The VML free energy update equations

To identify the update equations for the maximization of the VML free energy, we proceed as in Section S1.2. Because the main difference between the VB and VML framework is the parameterization of the error covariance matrix *V*_λ_ in terms of λ rather than *m*_λ_ and the vanishing of terms relating to the prior and variational distributions of λ, we can keep the discussion very concise.

The relevant subpart of 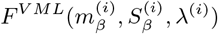 depending on *m_β_* is given by

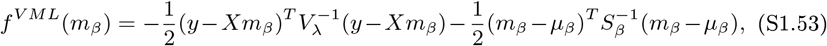

with gradient

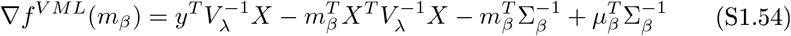

and ensuing update equation

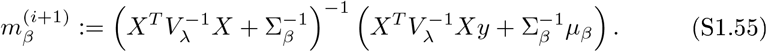

Likewise, the relevant subpart of 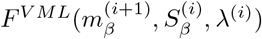 depending on *S_β_* is given by

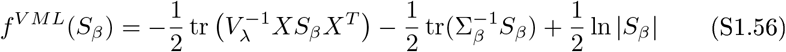

with gradient

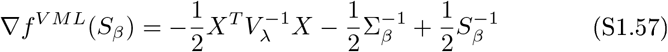

and the resulting update equation

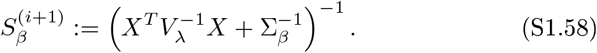

Finally, the relevant subpart of 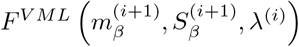 depending on λ is given by

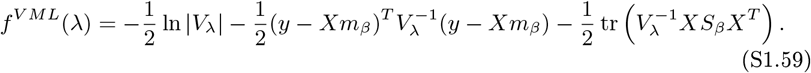

Here, in analogy to eqs. (S1.48) and (S1.49), the entries of 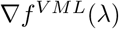 for the case of the two-component error covariance matrix of interest (eq. (S1.25)) evaluate to

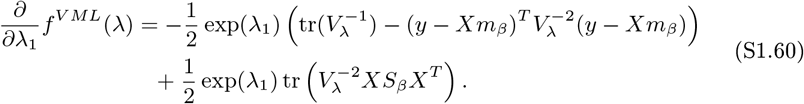

and

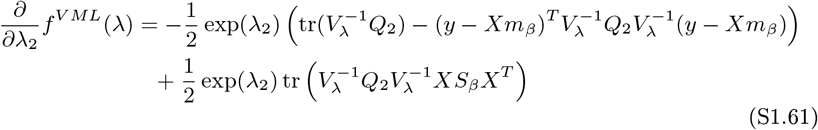

### S1.4 Restricted maximum likelihood

#### The ReML objective function as VML free energy

We first show that for the probabilistic model

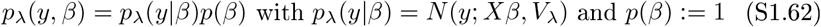

it holds that the VML free energy with variational distribution

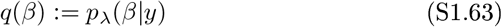

evaluates to the ReML objective function

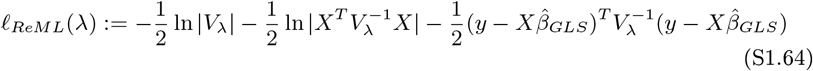

up to an additive constant, i.e.

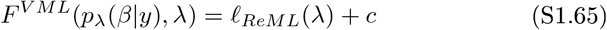

with

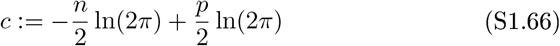

To this end, we first note that for the probabilistic model (S1.62) and with the definition of the GLS estimator

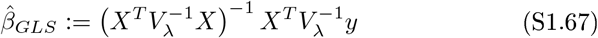

it holds that

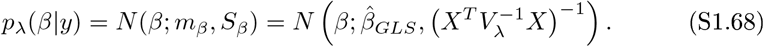

In brief, (S1.68) follows as a limiting case of the conditional properties of Gaussian distributions for the case of zero prior precision, i.e. the case of an improper prior *p(β)* = 1 (see e.g. Murphy (2012) for a more detailed discussion).

Evaluation of the VML free energy in the current scenario then yields

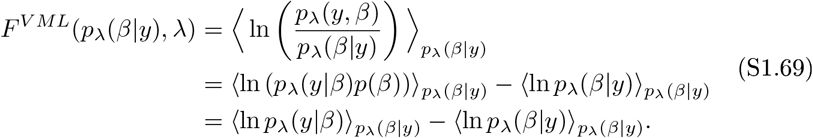

Evaluation of the first term on the right-hand side (S1.69) yields

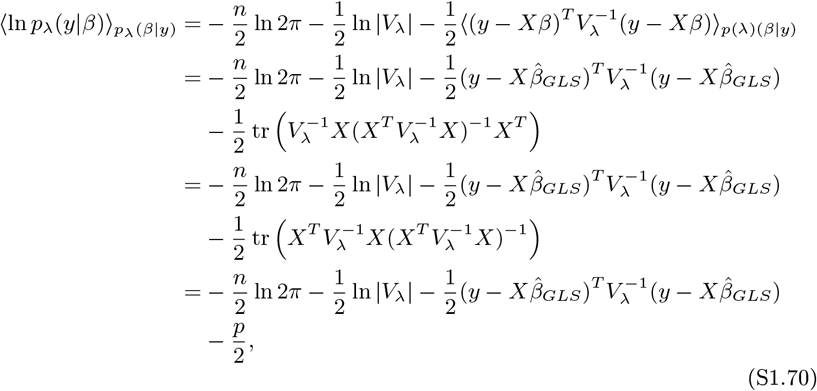

where the second equality follows with (S1.2). The third equality uses the invariance of the trace under cyclic permutations. The second term on the right hand of (S1.69) corresponds to the entropy of the distribution *_pλ_(β|y)* and thus evaluates to

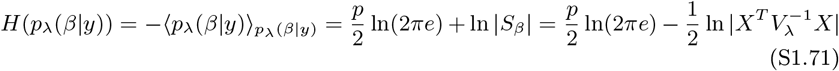

We thus have shown that

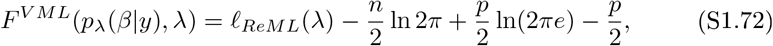

which concludes the derivation.

#### Evaluation of the ReML free energy function

To align the discussion of ReML with the previous discussions of VB and VML, we next define the ReML free energy function as the VML free energy evaluated for the probabilistic model (S1.62) at the exact posterior distribution *_pλ_(β|y)*, i.e.,

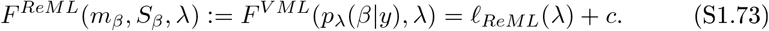

By noting that with (S1.68) the variational parameters are given by

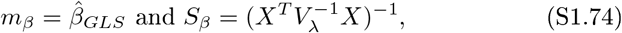

we can then rewrite the ReML free energy as in the main text:

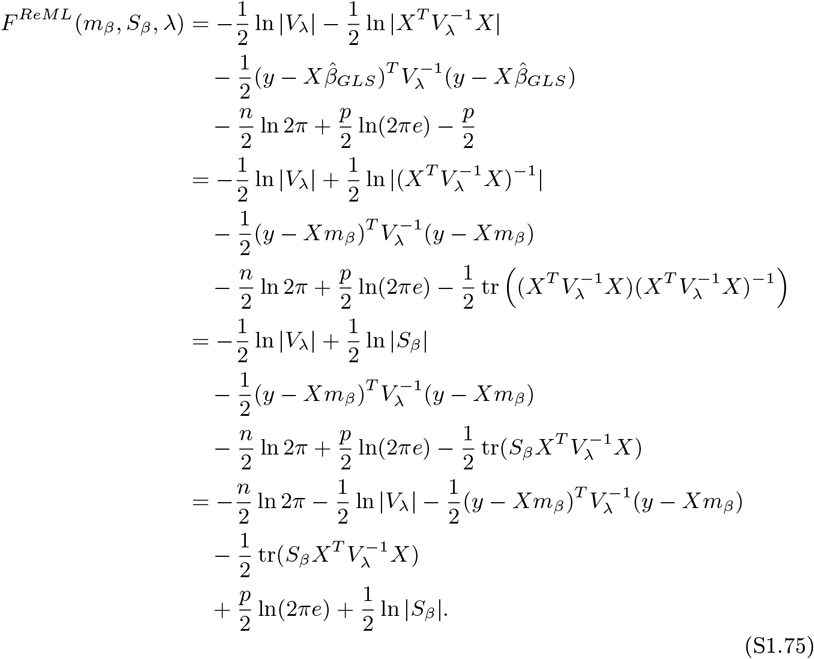

#### The ReML free energy update equations

Finally, we derive the update equations for the parameters *m_β_, S_β_*, and λ of the ReML free energy. Note that because the ReML objective function is identical to the ReML free energy up to an additive constant which is independent of these parameters, the resulting iterative algorithm also maximizes the ReML objective function.

The relevant subpart of 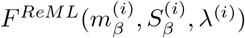 that depends on *m_β_* is given by, omitting iteration superscripts for ease of notation,

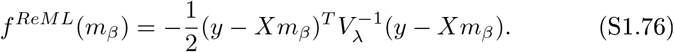

with gradient

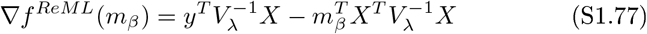

and ensuing update equation

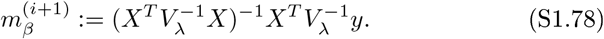

Unsurprisingly, this is the GLS estimator. Further, the relevant subpart of 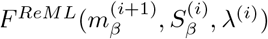 depending on *S_β_* is given by, again omitting iteration superscripts for ease of notation,

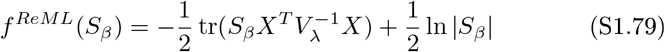

with gradient

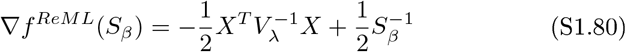

and ensuing update equation

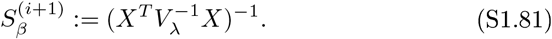

Finally, because the subpart of *F^ReML^* depending on λ is identical to the subpart of *F^VML^* depending on λ, the update procedure for *F^ReML^* with λ *F^VML^*.

### S1.5 Maximum likelihood

#### The ML free energy update equations

For the GLM, we have by definition

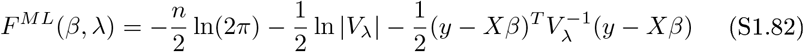

To derive parameter update equations, we consider the dependency of *F^ML^* on *β^(i)^* and λ^(*i*)^ in turn. The relevant subpart of *F^ML^*(*β^(i)^, λ^(i)^*) that depends on *β* is then given by, omitting iteration superscripts for ease of notation,

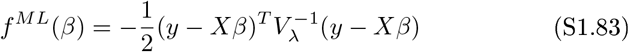

with gradient

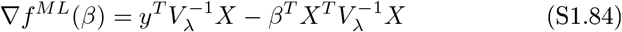

and ensuing update equation

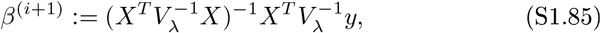

corresponding to the GLS estimator as in the case of ReML. The relevant subpart of *F^ML^*(*β^(i+1)^*, λ^(*i*)^) that depends on λ differs from the VML and ReML scenarios and is given by, again omitting iteration superscripts for ease of notation,

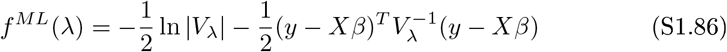

Here, in analogy to eqs. (S1.47), (S1.48), and (S1.49), the entries of 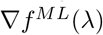 for the case of the two-component error covariance matrix of interest evaluate to

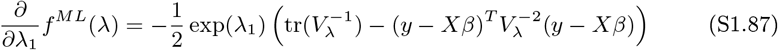

and

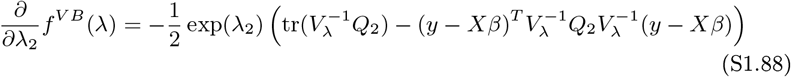

As they correspond to a disregard of prior information and posterior uncertainty about *β*, equations (S1.85), (S1.87) and (S1.88) can also be attained from the VML update equations (S1.55), (S1.60) and (S1.61) by setting 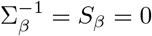.

## S2 Foundations of variational Bayes

In this section we formulate a probability-theoretic model of the probabilistic model considered in the main text in order to derive the VML and ML scenarios as special cases of VB. By “probability-theoretic” we mean a measure theory-based approach to probabilistic concepts, as prevalent in contemporary mathematics (e.g. Billingsley, 2012; Shao, 2003; Fristedt and Gray, 1997). This approach is rather uncommon in the neuroimaging and machine learning literature, where many application-oriented developments on VB have taken place (Blei et al., 2016). In the current context, it is necessitated by the fact that “point probability masses” cannot be represented by probability density functions. This implies that to derive VML and ML under VB requires a careful differentiation between those random variables whose distribution can and cannot be represented by probability density functions. This is afforded by the measure theory-based approach. We assume that the reader is familiar with the measure-theoretic viewpoint of probability theory, including Lebesgue integration. To establish notation and prepare some aspects of the discussion to follow, we provide a brief summary of key elements in Section S2.1. In Section S2.2 we then review a selection of entropy formulations which will be required for the formulation of VB and VML in probability-theoretic terms. Finally, in Section S2.3 we formulate the VB, VML, and ML scenarios is probability-theoretic terms and discuss their mutual relationships.

### S2.1 Preliminaries

#### Measurable, measure, and probability spaces

Our formulation rests on the concepts of *measurable, measure*, and *probability spaces.* A *measurable space* is a pair 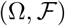, where Ω denotes a set and 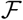 denotes a σ-field on Ω. An important measurable space in the following will be 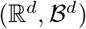, where 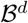 denotes the *d*-dimensional Borel σ-field on 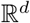. A *measure space* is a triple 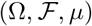, where *μ* denotes a measure, i.e. a mapping μ: 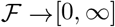 with properties

(Ml) 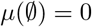, and

(M2) for every pairwise disjoint sequence 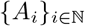 with 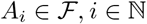 it holds that 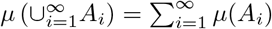.

An important measure space in the following will be 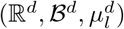, where 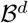 denotes the *d*-dimensional *Borel σ-field* and 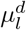 denotes the *d*-dimensional *Lebesgue measure*

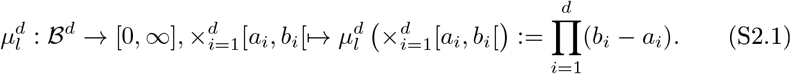

Please note that we do not use the more conventional notation λ^*d*^ for the Lebesgue measure to avoid confusion with the covariance component parameter vector λ. Similarly, a *probability space* is a triple 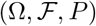, where *P* denotes a probability measure, i.e. a mapping 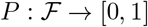, with propertie

(PI) 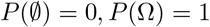, and

(P2) for every pairwise disjoint sequence 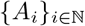 with 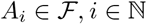 it holds that 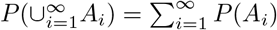.

An important probability space in the following will be 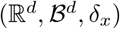, where *δ_x_* denotes the *Dirac measure*, defined as

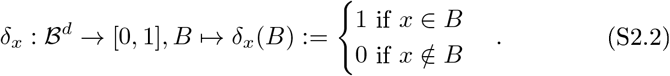

Of key importance in the derivations to follow is the fact that the Lebesgue integral of a measurable function 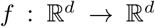 with respect to the Dirac measure *δ_x_* is readily evaluated as (e.g. Lieb and Loss, 2001)

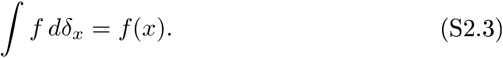

#### Random variables and distributions

Let 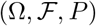 and 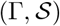 denote a probability space and a measurable space, respectively. A *random variable* is a function

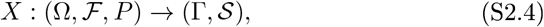

which is measurable, i.e. for which

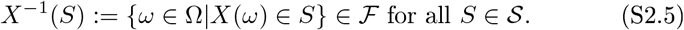

A random variable induces a probability measure

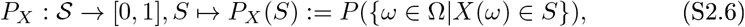

on 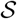. The probability measure *P_X_* is referred to as the *distribution* of the random variable *X* and renders 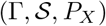 a probability space (e.g. Fristedt and Gray, 1997, Chapter 2).

#### Probability density functions

For a measure space 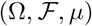 any quasiintegrable function 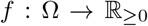 is a *density function* and defines a measure 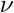 on 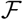 by means of its Lebesgue integral for 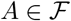, i.e.

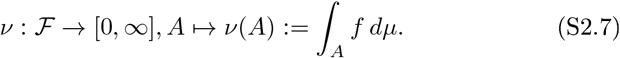

We say that “*v* is a measure with density function *f* with respect to the measure *μ*” and write 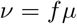 for short. Recall that a 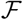-measurable function 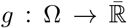 is integrable with respect to 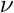, if the function product *g* · *f* is integrable with respect to *μ*, and that in this case

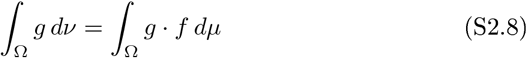

(e.g. Billingsley, 2012, Theorem 16.11). As noted above, we will be primarily concerned with the measure space 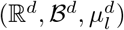, where 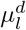 denotes the Lebesgue measure on 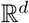. In this case, if the Lebesgue integral of *f* with respect to 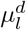 equals 1, *f* is referred to as *probability density function*. Furthermore, in this case we have for 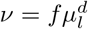

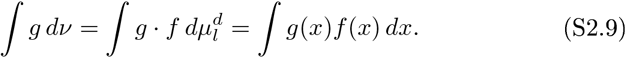

Crucially, this implies that one can evaluate Lebesgue integrals using Riemann integration as done throughout Section S1 (right-hand side of (S2.9), for details see e.g. Schmidt (2011), Chapter 9).

We further require the notion of conditional probability density functions. For a random variable (*X, Y*) on a product measure space 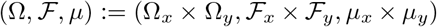 with joint probability density function 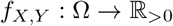 with respect to *μ*, the conditional probability density function of *X* given *Y = y* is defined as

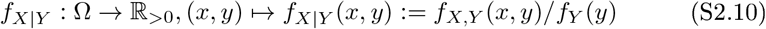

where

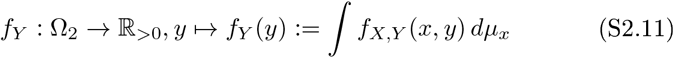

is the marginal probability density function of *Y* with respect to *μ_y_* (e.g. Shao, 2003, Chapter 1.4).

#### Discrete, continuous, and mixed random vectors

Because we are considering multivariate random entities in the application of VB, VML, and ML to the GLM, we also require the notion of *random vectors* as the multivariate extension of random variables. More specifically, we require the concepts of *discrete, continuous* and *mixed random vectors*, which we introduce in the following.

Let 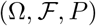 be a probability space. A *d*-dimensional *discrete random vector* is a function

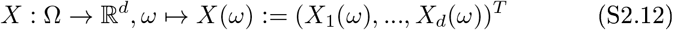

whose range space or *alphabet* (Gray, 2011; Cover and Thomas, 2012)

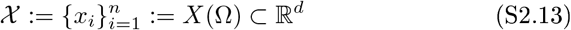

is finite. A discrete random vector has an associated *probability mass function p_X_* given by

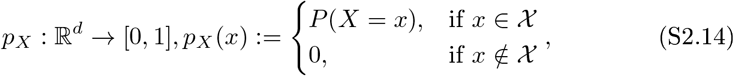

with the notational convention

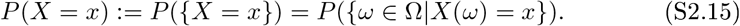

A special discrete random vector required in the following is a *constant random vector*, which for a fixed 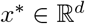 we write as

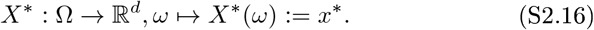

In this case the alphabet 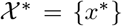 of X* comprises a single element, the associated probability mass of which is given by

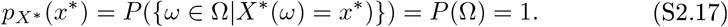

Analogously to a random variable, a random vector 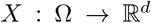 induces a probability measure *P_X_* on the measurable space 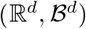. Notably, the induced probability measures of a constant random vector is the Dirac measure (e.g. Bauer, 1991, p. 25), i.e. with (S2.2) and (S2.16)

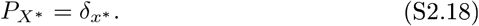

A *d*-dimensional *continuous random vector* is a function *Y* of the form (S2.12) whose induced probability measure *P_Y_* on 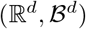 is absolutely continuous with respect to Lebesgue measure and can thus be represented by a probability density function 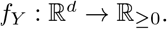.

Finally, we construct the concept of a *mixed random vector* as follows. Set 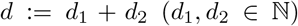, let 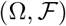 be a measurable space, and let 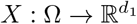 be a discrete random vector, which induces a distribution *P_X_* on 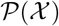. Let *P_Y|X_* be a Markov kernel from 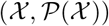 to 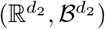, i.e. *P_Y|X_* is a mapping

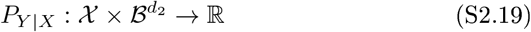

with the properties

- 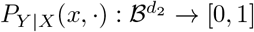 is a probability measure on 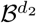 for every 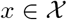,
- 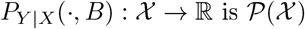-measurable for every 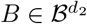.

Then *P_X,Y_ = P_X_P_Y|X_* is a probability measure on 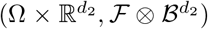 (e.g. Shao, 2003, 1.4.3). Assume in addition that

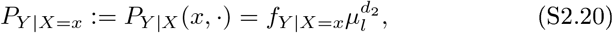

i.e.

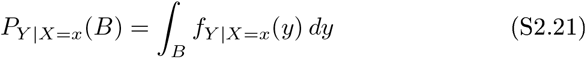

for probability density functions *f_Y|X=x_* with 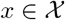. Let *Y* denote the identity-mapping on 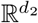 and define *Z* := (*X, Y*). Then *Z* is a *d*-dimensional random vector on 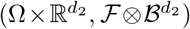 with marginal distributions having the properties

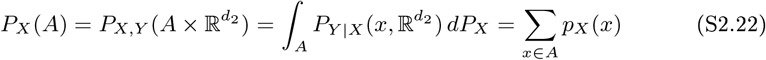

for all 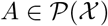 and

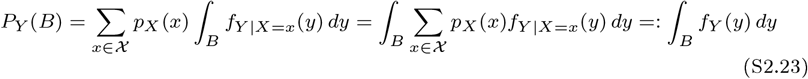

for all 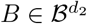. In other words, *X* is (by definition) a discrete *d*_1_-dimensional random vector (which we call the *discrete component* of *Z*) and *Y* is (by construction) a continuous *d*_2_-dimensional random vector (which we call the *continuous component* of *Z*). We call *Z* a *d*-dimensional *mixed random vector*.

Note that if we set *P_Y|X_*(*x*, ·) := *P_Y_* for every 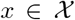 with a probability measure *P_Y_* on 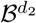, then *P_X,Y_ = P_X_P_Y_* is a product measure and the random vectors *X* and *Y* are independent. Vice versa, assuming independent *d*_1_ - and *d*_2_-dimensional random vectors *X* and *Y*, respectively, we can use the construction above to construct a *d*-dimensional mixed random vector with discrete and continuous components whose marginal distributions are independent.

### S2.2 Entropies of distributions of random vectors

#### Entropy of the distributions of a discrete random vector

Following (Gray, 2011, Chapter 3), we define the entropy of the distribution *P_X_* of a discrete random vector *X* with alphabet 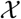 as

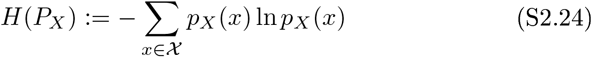

with the convention 0 ln 0 := 0. For later reference we note that the the entropy of the distribution of a constant random vector of the form (S2.16) is zero, because in this case the defining sum (S2.24) comprises a single term which evaluates to zero:

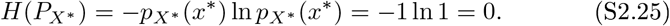

#### Entropy of the distribution of a continuous random vector

We define the entropy of the distribution *P_Y_* of a continuous random vector *Y* as its *differential entropy* (e.g. Cover and Thomas, 2012, Chapter 8), i.e. we set

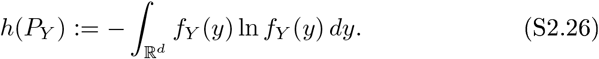

#### Entropy of the distribution of a mixed random vector

Finally, following (Nair et al., 2006), we define the entropy of the distribution *P_Z_* of a mixed random vector *Z = (X, Y)* with the property

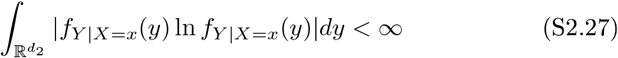

for all 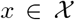 by

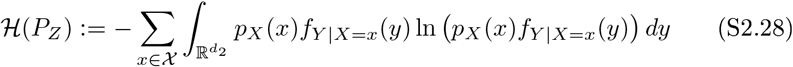

Note that we can rewrite this definition as

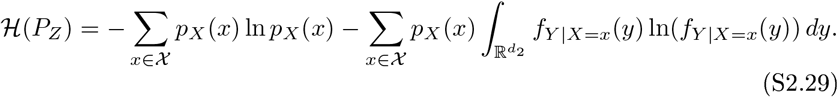

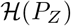 thus comprises the sum of the entropy of the marginal distribution of the discrete component of *Z* and a convex combination of the differential entropies of the conditional distributions of the continuous components.

In particular, for the case of independent discrete and continuous components, i.e. *f_Y|X=x_ := f_Y_*, we obtain

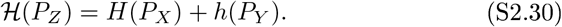

More generally, if the components *X* and *Y* are independent, and *Y* may be either continuous or discrete (rendering *Z* a discrete random vector), we write

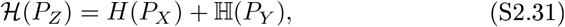

where 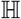 denotes the entropy of the continuous or discrete component *Y*.

### S2.3 Probability-theoretic variational Bayes

Based on the concepts reviewed in Sections S2.1 and S2.2, we are now in the position to formulate the VB, VML, and ML scenarios discussed in the main text in probability-theoretic terms and to delineate their relationship. We proceed as follows. First, we reformulate the free energy functions *F^V B^, F^V M L^* and *F^M L^* introduced in the main text in probability-theoretic terms. To distinguish these functions from their counterparts in the main text, they will be denoted by 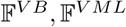, and 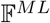, respectively. Note that in general, the free energy functions depend on both the realizations of the observed data random variables as well as the distributions (or values in the VML and ML case) of the unobserved parameter random variables (or non-random variables in the VML and ML case). However, in analogy to classical likelihood functions (e.g. Shao, 2003, Chapter 4.4), we conceive of the free energy functions as functions of entities related to the parameter (random) variables only. Intuitively, this corresponds to assumption of a given and fixed data observation - the common scenario in experimental applications of the approaches. Second, upon reformulating the VB, VML, and ML scenarios in probability-theoretic terms, we relate these new formulations to the definitions of the variational free energies in the main text and show their consistency. Finally, we conclude with a theorem on the relationshiop between VB, VML, and ML.

#### Variational Bayes

To express the VB scenario of the main text in probability-theoretic terms, we set 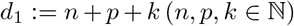, consider a probability space 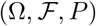 and the measurable space 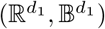 and define the continuous random vector

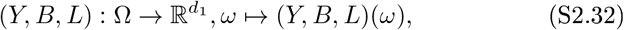

which induces a probability measure *P_Y,B,L_* on 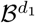. We thus obtain the probability space 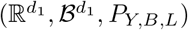. Because (*Y, B, L*) is a continuous random vector, *P_Y,B,L_* can be represented by a probability density function

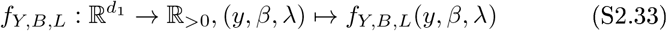

with respect to Lebesgue measure 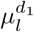 on 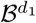. Note that *f_Y,B,L_* is denoted by *p(y, β, λ)* in the main text.

To define the variational Bayes free energy function 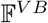, we first consider a random vector

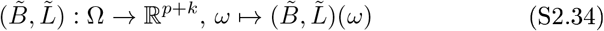

whose components are the independent random vectors 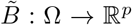 and 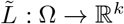. This implies that the induced distribution 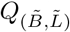 on the measurable space 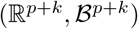 of 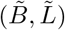 factorizes, i.e.

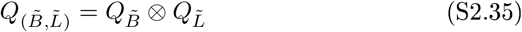

where 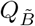 and 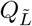 denote the marginal distribution on 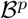 and 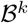 induced by 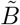 and 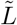, respectively (e.g. Fristedt and Gray, 1997, Chapter 9). We hence write 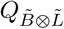 for 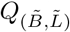. Let 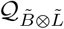 denote the set of all such distributions. For a fixed 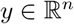 we then define

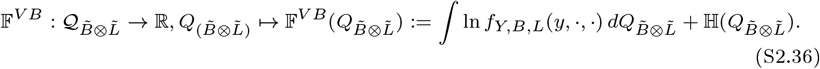

Here the symbol 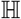 denotes the entropy of the distribution 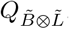, the evaluation of which depends on the type of random vectors 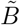 and 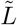, as discussed in Section S2.2.

#### Variational maximum likelihood

In analogy to the above, we set 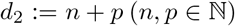, consider a probability space 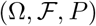 and the measurable space 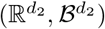, and define the continuous random vector

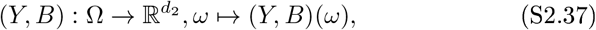

which induces the probability measure *P_Y,B_* on 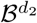. We thus obtain the probability space 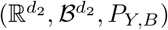. As in the main text, we assume that *P_Y,B_* is represented by a parameter-dependent probability density function

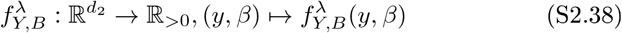

with respect to Lebesgue measure on 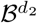. Note that 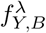 is denoted by *p_λ_*(*y, β*) in the main text.

To define the variational maximum likelihood free energy function 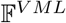, we first consider a random vector 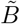

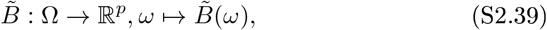

which induces a probability measure 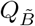 on the measurable space 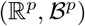. Let 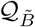 denote the set of all such induced probability measures. For a fixed 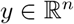 we then define

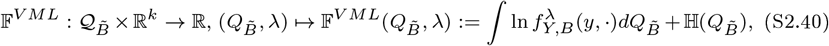

where as above the symbol 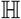 denotes the entropy of the distribution 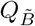, the evaluation of which depends on the type of random vector 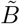 as discussed in Section S2.2.

#### Maximum likelihood

Finally, to express the maximum likelihood scenario in probability-theoretic terms, we consider a probability space 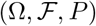, the measurable space 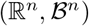 and define the continuous random vector

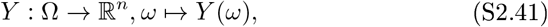

which induces the probability measure *P_Y_* on 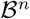 and hence the probability space 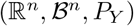. As in the main text, we assume that *P_Y_* is represented by a parameter-dependent probability density function

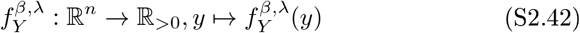

with respect to Lebesgue measure on 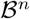. Note that 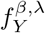 is denoted by *_pβ,λ_(y)* in the main text. We define the maximum likelihood free energy function 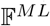 as the standard log-likelihood function of the maximum likelihood scenario, i.e. for fixed 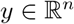, we set

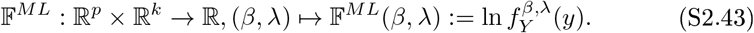

Note that we have 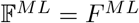 by definition.

This concludes the probability-theoretic formulations of the VB, VML, and ML, scenarios. We next show the consistency of the free energy functions defined in this section with the definitions of the free energy functions considered in the main text in form of the following lemma:

##### Lemma (Consistency of free energy function definitions).

*The definitions of the variational free energy functions* 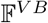 *and F^V B^, as well as* 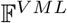 *and F^V M L^ are consistent. More specifically*,

(LI) *if in the definition of the variational Bayes free energy* 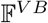 (S2.36) 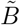 *and* 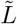 *are continuous random vectors represented, by probability density functions* 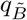 *and* 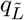 *with respect to Lebesgue measures* 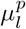 *and* 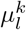, *respectively, then the definitions of* 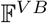 *and F^V B^ are equivalent, and*
(L2) *if in the definition of variational maximum likelihood free energy* 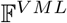 (S2.40) 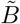 *is a continuous random vector represented, by a probability density function* 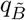 *with respect to Lebesgue measure, then the definitions of* 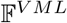 *and F^V M L^ are equivalent*.

##### Proof of (LI).

Consider the definitions

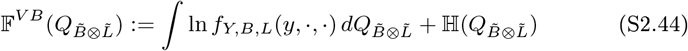

and, under the mean-eld approximation,

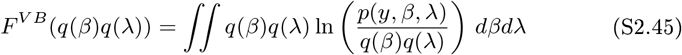

Note that the integral in (S2.44) denotes a Lebesgue integral with respect to the probability measure 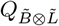, while the integral in (S2.45) denotes a (double) Riemann integral. If 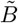 and 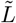 are continuous random vectors with associated probability density functions 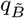 and 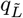, then with (S2.9) and the notational conventions of the main text, we have

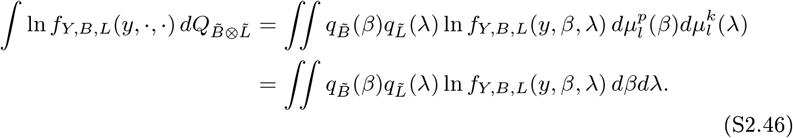

Further, because 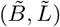 is a continuous random vector, (S2.26) applies for the entropy of 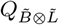, and thus

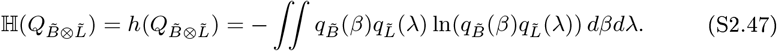

Hence, (L1) follows with the linearity of the (Riemann) integral and omission of the subscripts 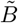 and 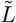 in the denotation of the probability density functions 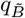 and 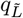.

##### Proof of (L2).

Consider the definitions

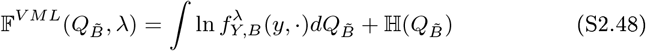

and

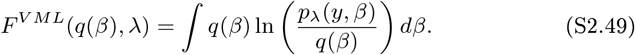

Note that the integral in (S2.48) denotes a Lebesgue integral with respect to the probability measure 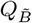, while the integral in (S2.49) denotes a Riemann integral. If 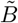 is a continuous random vector with associated density function 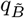 with respect to Lebesgue measure, then with (S2.9) and the notational conventions of the main text, we have

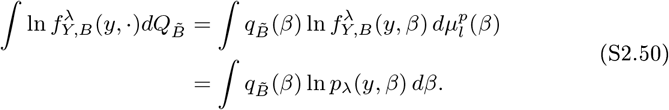

Further, because 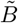 is a continuous random vector, (S2.26) applies for the entropy of 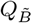, and thus

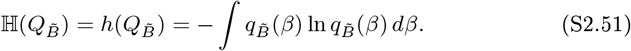

Hence, (L2) follows with the linearity of the (Riemann) integral and omission of the subscript 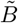 in the denotation of the probability density function 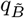.

Finally, we provide the key result of this section:

#### Theorem (Relationship of VB, VML, and ML).

*Variational maximum likelihood and maximum likelihood are special cases of variational Bayes. More specifically*:

(Tl) *For a constant marginal density f_L_(λ) := 1 and the constant random vector*

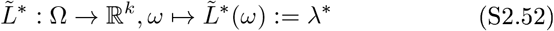

*it holds that*

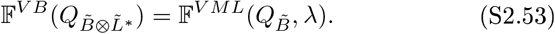
(T2) *For a constant marginal density f_B_ (β) := 1 and the constant random vector*

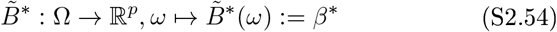

*it holds that*

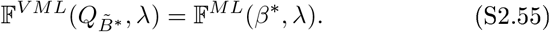

##### Proof of (Tl)

We first note that because 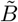 and 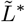 are independent random vectors and because 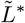 is a constant random vector, we have with (S2.30), (S2.31), and (S2.25)

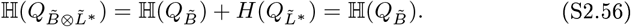

Second, with (S2.18) we have 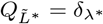. Hence, with Fubini’s theorem,

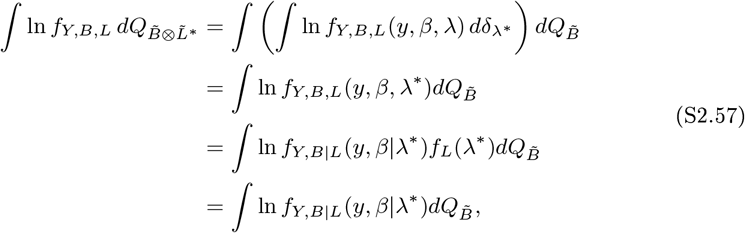

where the last equality follows with *f_L_*(λ*) = 1. Notationally identifying the probability density function

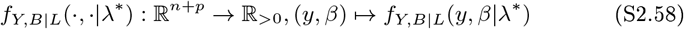

with the probability density function 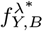 and omission of the asterisk superscript then completes the proof.

##### Proof of (T2)

We first note that because 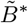 is a constant random vector, we have with (S2.25) 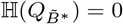. Second, with (S2.18) we have 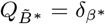. Hence,

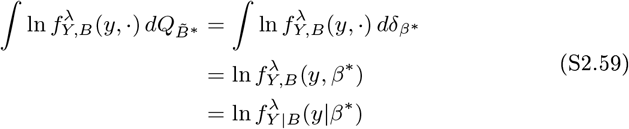

**Figure S1:**
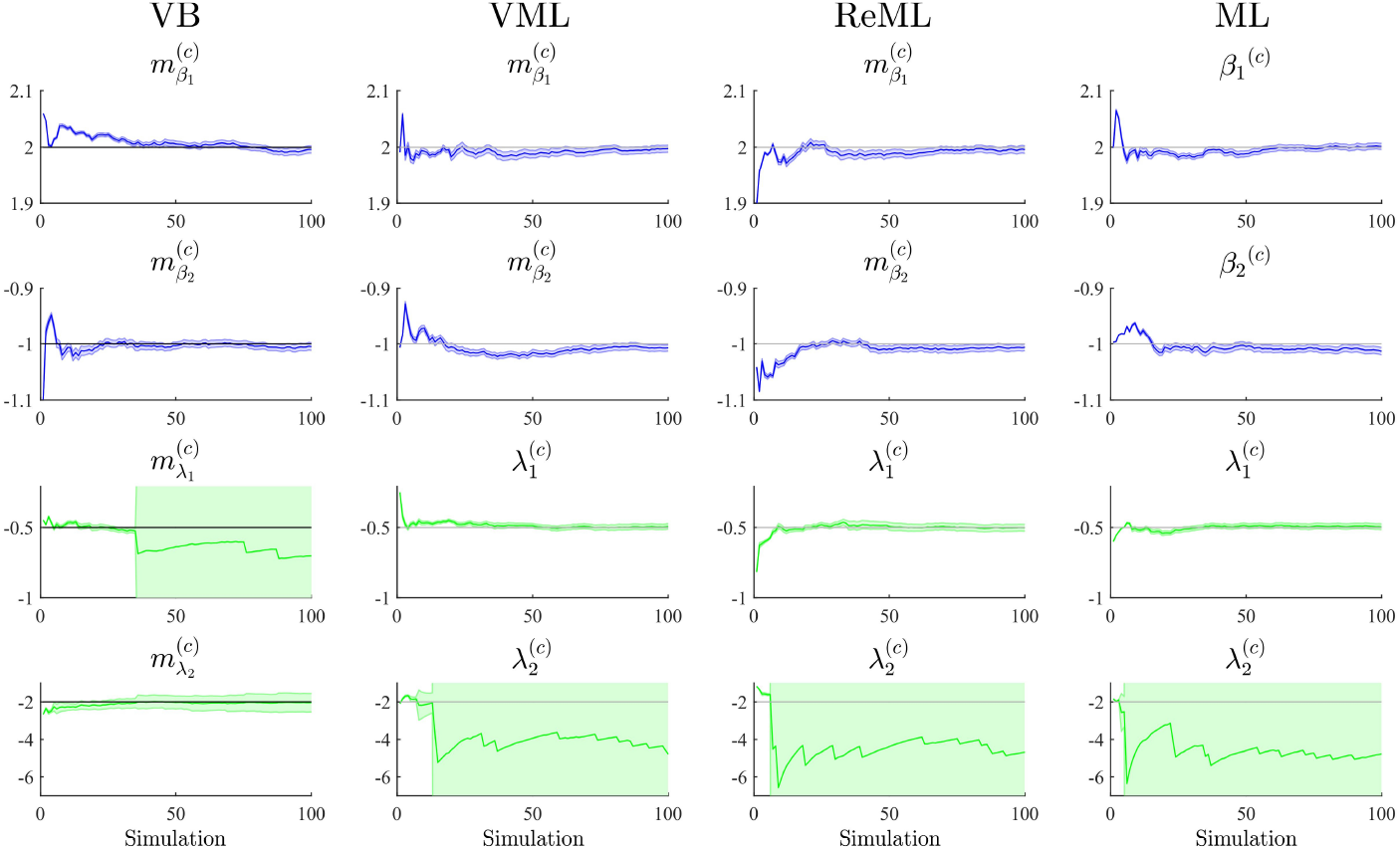
The panels along the figure’s columns depict the cumulative averages (blue/green lines), cumulative variances (blue/green shaded areas), and true, but unknown, parameter values (grey lines) for VB, VML, ReML, and ML estimation. Parameter estimates relating to the effect sizes *β* are visualized in blue, parameter estimates relating to the covariance components λ are visualized in green. The panels along the figure’s rows depict the parameter recovery performance for the subcomponents of the effect size parameters (row 1 and 2) and covariance component parameters (row 3 and 4), respectively. As opposed to the data shown in the main text, the covariance component parameter estimates are not corrected for outliers. For implementational details, please see 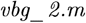.

where the latter equality follows with *f_B_ (β*)* = 1. Notationally identifying the probability density function

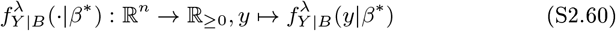

with the probability density function 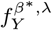 and omission of the asterisk superscript then completes the proof.

## S3 Cumulative averages without outlier removal

In Section 3.1 of the main text, we present a parameter recovery simulation for which we removed 15-20 % of outliers in the estimation of the covariance component parameters. The same results without outlier removal are depicted in Figure S1. As evident from the Figure, outliers aect all four estimation techniques, but primarily one of the covariance components. Retaining the outliers results in negative estimation bias estimates and an increase of the estimation variance estimate.

**Figure S2:**
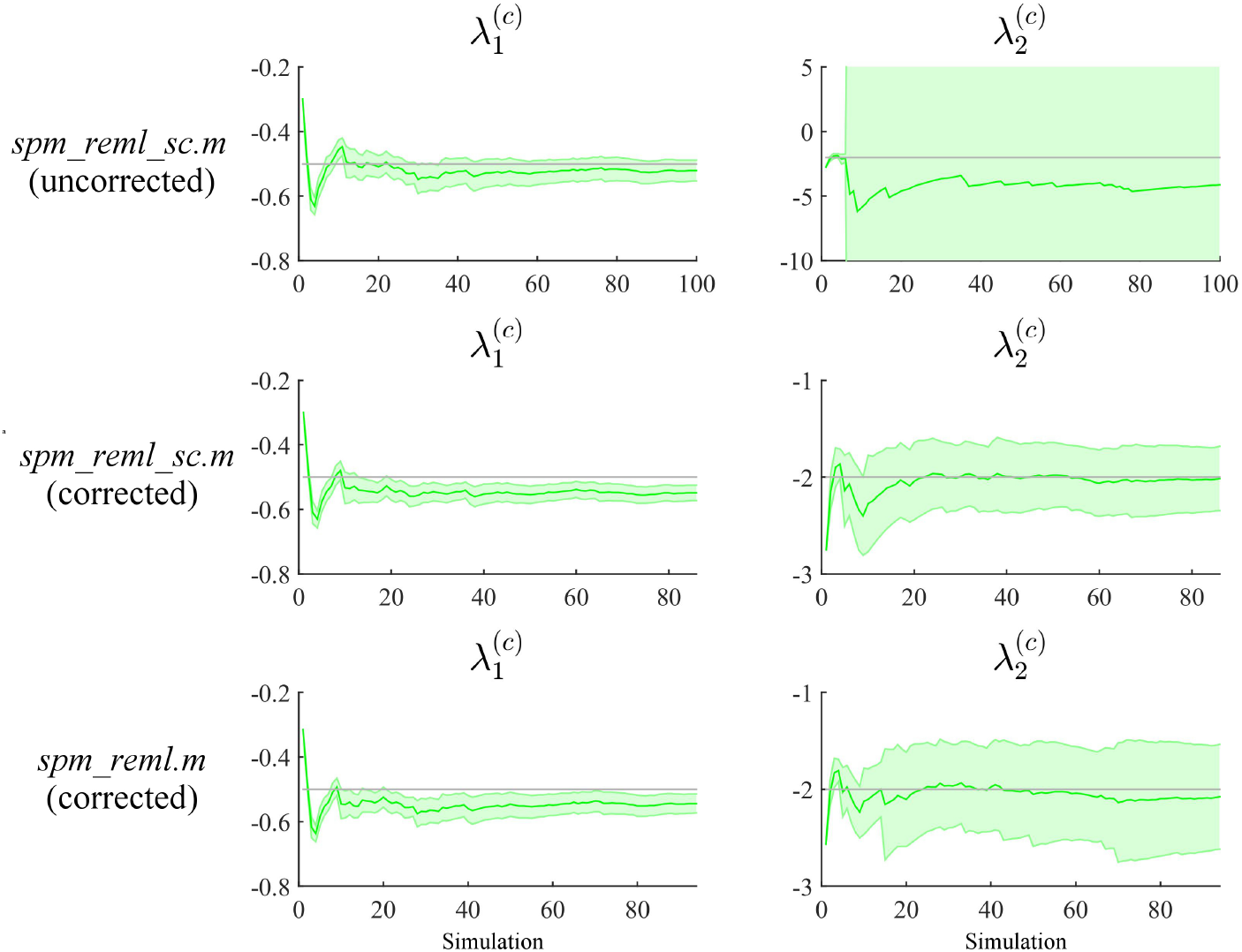
Parameter recovery for SPM12-based covariance component parameter estimation. The panels along the figure’s columns depict the cumulative averages (gree line), cumulative variances (green shaded area), and true, but unknown, parameter values (grey) for the first and second covariance component parameters λ_1_ and λ_2_, respectively. The panels along the figure’s rows depict these quantities for the two implementations of covariance component parameter estimation in SPM12 as indicated on the right, and without and with a correction for outliers as indicated. For implementational details, please see 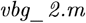.

## S4 SPM12 ReML estimation

In the parameter recovery assessment of our VB, VML, ReML, and ML implementation, we found that the covariance component parameter estimation fails in a significant number of cases. To investigate whether this behaviour is specific to our implementation, we performed the same analyses using the covariance component parameter estimation functions *spm_reml_sc.m* (Version 4805) and *spm_reml.m* (Version 5223) of the SPM12 distribution. These functions perform a Fisher scoring ascent on the ReML objective function to identify maximum-a-posteriori covariance component parameter estimates, probably documented best in (Friston et al., 2002). The function *spm_reml_sc.m* uses weakly informative log normal priors to ensure the positivity of the covariance component parameter estimates, while the *spm_reml.m* function, which is called by SPM12 central *spm_spm.m* function, does not.

We visualize the results in Figure S2. The panel columns of this figure refer to the two covariance component parameter estimates and the panel rows refer to the different SPM12 functions. In the first row, we visualize the cumulative average and variances of the respective parameter estimates based on the *spm_reml_sc.m* function without the removal of outliers. The performance for λ_1_ is acceptable, but for the estimation of λ_2_ outliers from approximately the 10th simulation on bias the cumulative average significantly away from the true, but unknown, parameter value and strongly amplify the cumulative variance. This is similar to the behaviour we detected in our implementation which led us to remove these outliers automatically (Grubbs, 1969). The second row of Figure S2 depicts the parameter recovery performance for *spm_reml_sc.m* after removal of appoximately 15% of outliers. This results in similar performance as in our implementation. Finally, the last row of Figure SS2 depicts the parameter recovery performance for the *spm_reml.m* function. Because *spm_reml.m* can return negative covariance components and because the SPM12 procedures assume a covariance structure of the form 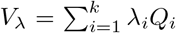 and not of the form 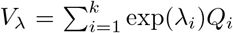 as in our implementation, the necessary log transformation of the returned parameter estimates here can result in undefined results. In the data shown, these undefined results have been removed, again rendering the resulting cumulative averages and variances within reasonable bounds of the true, but unknown, parameter values.

In summary, we conclude that the numerical optimization problems that we encountered for the estimation of covariance components based on our implementation of the VB, VML, ReML, and ML estimation techniques are not an uncommon phenomenon in the analysis of nouroimaging data.

## S5 Model recovery free energy contributions

To understand the observed pattern of average free energies in Figure 7 in further detail, we tabulated the sum terms of each free energy function (Table S1) and visualize the average term contributions to the overall average free energy in Figure S3. We omit from visualization the first term, which is identical for all free energy functions and evaluates to T1 = −367.58. Of the remaining terms, the largest contributions are provided by T3 and T2, reecting the residual sum of squares and the log determinant of the estimated data covariance matrix, respectively (Figure S3A). The remaining terms T4 - T17, as far as they exist for each free energy function, make smaller contributions (Figure S3B). Notably, the residual sum of squares is virtually identical over all pairings of data generating and data analysis model. This reects the fact that the two-regressor model MA2 can readily capture the data pattern of the single-regressor model MG1 by estimating β_2_ to be approximately zero. The average free energy differences for the two data analysis models in case of MG2 thus appear to be primarily accounted for by the different contributions of T2. It is likely that these differences result from the erroneous allocation of data variance under model MG2 to the covariance components of model MA1. The more subtle differences between the average free energies for MA1 and MA2 in the case of MG1 on the other hand, seem to arise from two factors: firstly, a slight overestimation of the covariance component parameters of MA2 in case of MG1, leading to a persistence of the lower average free energy values in the case of ML estimation, and secondly, from additional contributions of T4 - T16 for VB, VML, and ReML, as evident from the more negative sums (Σ) of these terms. In summary, for the current data generating and data analysis model comparison, both covariance component overestimation and the free energy model complexity terms T4 - 16 appear to contribute to the identiability of the true, but unknown, model structure.

**Table S1:**
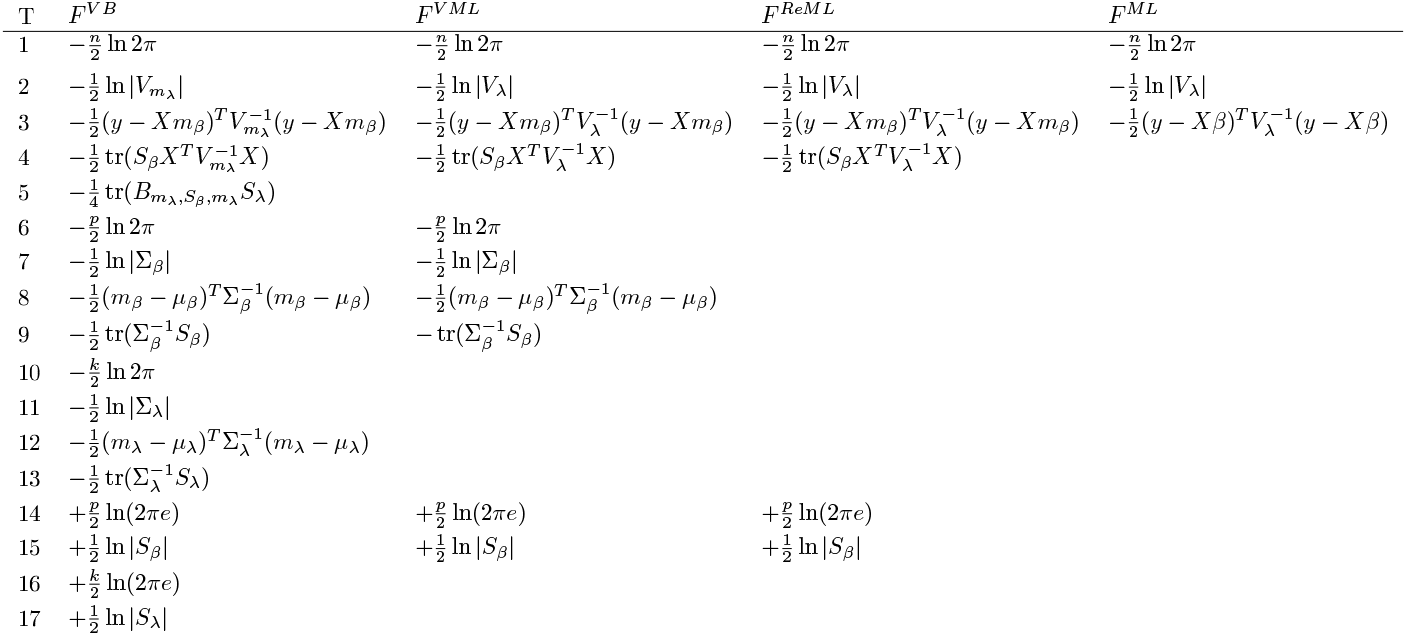
Free energy sum terms. Note that T1 is identical over all free energy functions, T2 is the negative log determinant of the estimated data covariance matrix, and T3 corresponds to the residual sum of squares. The remaining terms, if they exist relate to the prior and posterior uncertainties over model parameters and are commonly referred to as “model complexity” terms.

**Figure S3:**
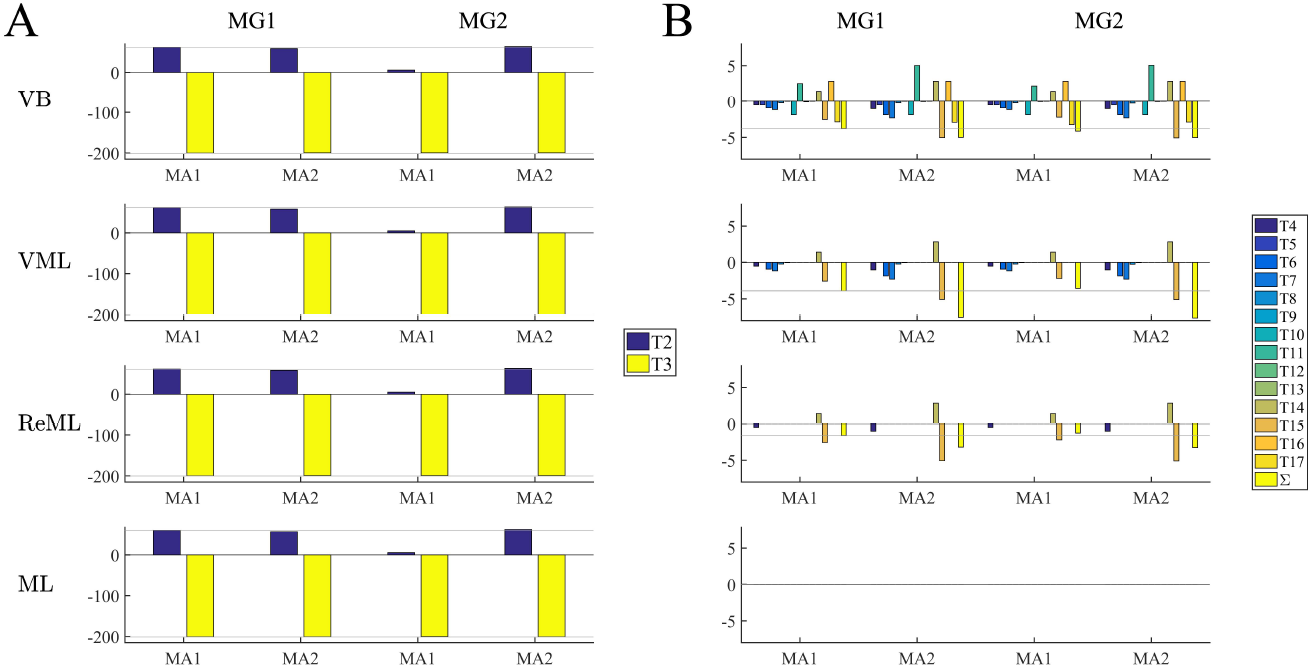
Average free energy term contributions. The figure depicts a decomposition of the average free energy values depicted in the main text Figure 7 according to the terms tabulated in Table S1. Panel A displays the largest contributions, afforded by terms T2 and T3, Panel B displays the remaining term contributions, and the sum (Σ) of these remaining distributions. Note the difference in scale between Panels A and B. Panel rows refer to the four estimation techniques. In each subpanel, the left two bar groups refer to data generated by MG1 analyzed with data analysis models MA1 and MA2, and the right two bar groups refer to data generated by MG2 analyzed with MA1 and MA2. For visual comparison, thin grey lines corresponding to the values obtained under the MG1/MA1 combination are included. For implementational details, please see 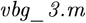.

